# Double Periodicity of the AnkyrinG-Associated Complex in the Axon Initial Segment

**DOI:** 10.64898/2026.04.25.720723

**Authors:** Gülberk Bayraktar, Josephine K Dannersø, Sean DS Hansen, Lisbeth Schmidt Laursen, U. Valentin Nägerl, Poul Nissen

**Author notes:** These authors contributed equally to the study (Gülberk Bayraktar, Josephine K Dannersø).

## Abstract

The axon initial segment (AIS), situated within the first 20-60 µm of the axon, is essential for action potential generation and maintenance of axonal identity. Its structure relies on the beta (β)-IV-spectrin/AnkyrinG (AnkG) scaffold arranged periodically underneath the plasma membrane, harbouring diverse membrane proteins. Although a ∼190-nm cytoskeletal periodic organization is well established, the precise stoichiometry and spatial arrangement of AIS proteins within the ∼190-nm spatial period remain rudimentary, mostly for lack of sufficient spatial resolution and labelling efficiency. Here, using expansion microscopy and cryo-electron tomography, which overcome these technical limitations, we present data on the organization of the AnkG-associated complex within the ∼190-nm spatial period. We demonstrate that exactly two AnkG molecules with their C-termini separated by ∼80 nm are situated within each period. By contrast, the AnkG-associated cell-adhesion protein neurofascin-186 appears in clusters of varying sizes that are consistent with the periodic organisation of AnkG pairs, yet suggest a more complex molecular arrangement between the two molecules. Altogether, our novel approach provides new insights into AIS molecular organisation and protein stoichiometry.

## Introduction

The axon initial segment (AIS) is a highly specialized axonal region (20-60 µm) that separates the somatodendritic compartment from the axon and plays a key role in maintaining neuronal polarity^1,2^. The AIS is characterized by a high number of voltage-gated sodium (Na_v_) and potassium channels (K_v_) with unique properties (e.g. low voltage threshold) that facilitate the generation of action potentials (AP), making it a unique site that transforms somatodendritic electrical activity into a sequence of APs^3–7^. The AIS ultrastructure is characterized by a membrane-associated and periodic scaffold (MPS) that establishes a regular nanoscale architecture as shown by super-resolution fluorescence microscopy and platinum replica electron microscopy^8–11^. Spectrin tetramers, formed by end-to-end association of α-II-spectrin/β-IV-spectrin heterodimers, span ∼190 nm and are linked by circumferential actin rings at each tetramer junction (Fig. 1a)^8,12,13^.

**Figure 1.**
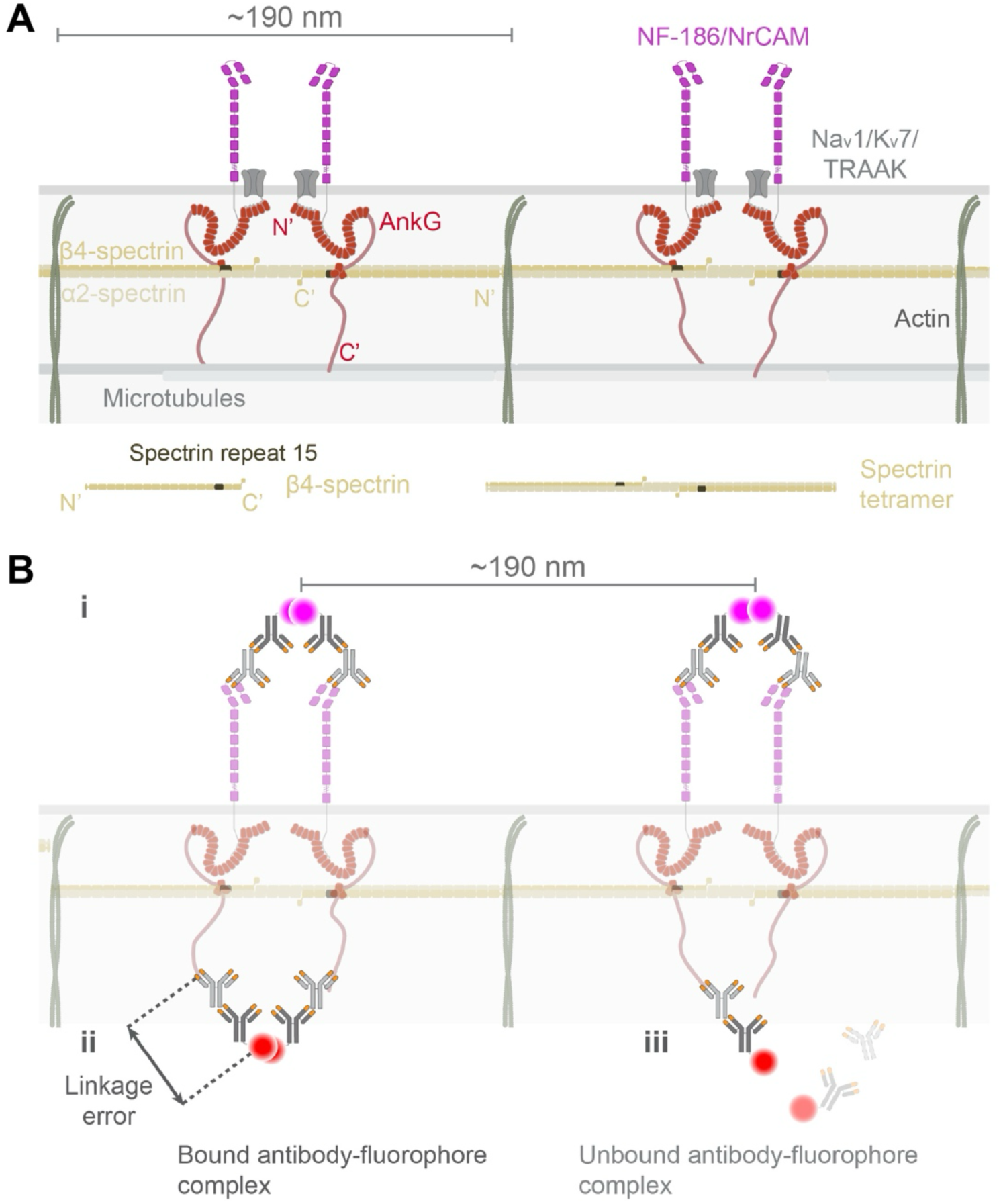
Nanoscale AIS Architecture and limitations of antibody-based super-resolution fluorescence microscopy in resolving it. **A,** The AIS is organised into repeating MPS units built on ∼190 nm long α-II/β-IV-spectrin tetramers. These spectrin tetramers are intercalated by actin rings and decorated with AnkG. β-IV-spectrin harbors interaction with AnkG at spectrin repeat 15, allowing up to two AnkG molecules per MPS unit. AnkG anchors membrane proteins including Na_v_1.2 and Na_v_1.6, K_v_7.2 and K_v_7.3, TRAAK, and the cell-adhesion molecules NF-186 and NrCAM to the spectrin scaffold, forming the AnkG-associated complex organised every ∼190 nm along the AIS. **B,** In current super-resolution microscopy, detected fluorescence does not fully reflect the underlying nanoscale architecture due to limitations imposed by probe geometry and the dense ultrastructure. **(i)** Closely spaced proteins cannot be reliably distinguished because the large antibody-fluorophore complex (∼20 nm) limits effective spatial precision, causing overlapping point spread functions (PSFs) that obscure neighbouring structures and true molecular distances. **(ii)** The size of antibody-fluorophore complex displaces the fluorophore from the target epitope, introducing linkage error and positional uncertainty. **(iii)** Reduced epitope accessibility and incomplete labelling can bias detection and hinder faithful reconstruction of the underlying molecular organisation.

Ankyrin proteins associate with spectrins to form cytoskeletal meshes in various cell types, which has been well studied in erythrocytes^14,15^. They are composed of a highly conserved N-terminal region featuring a membrane-binding domain (MBD), spectrin-binding domain (SBD), death domain and an intrinsically disordered C-terminal tail of varying lengths in different isoforms^14^. Ankyrins can bind multiple proteins simultaneously through the characteristic MBD harbouring 24 ANK repeats^16–18^.

The giant 480-kDa AnkyrinG (AnkG) is the dominant isoform in the AIS of neurons, where it plays a pivotal role in its organisation^19,20^. AnkG links a number of membrane proteins such as the cell-adhesion proteins Neurofascin-186 (NF-186) and NrCAM, Na_v_ isoforms 1.2 and 1.6, K_v_ isoforms 7.2 and 7.3, and the two-pore domain potassium channel TRAAK to the β-IV-spectrins (Fig. 1A)^21–26^. AnkG has been shown to bind to the spectrin repeat-15 of β-IV-spectrin^27,28^, resulting in two binding sites per spectrin tetramer. Altogether, the molecular organisation of spectrins and AnkG underlie the capacity of the AIS for dense packing of many proteins. Methodological advances in spatial resolution have provided new insights into this dense ultrastructure and its function.

Although two AnkG molecules are known to bind β-IV-spectrins within the MPS, direct visualization of such pairs has remained elusive. NF-186 is likewise predicted to occur in pairs associated with the MBDs of two AnkG molecules along the MPS. Despite the improvement in spatial resolution achieved by super-resolution fluorescence microscopy, the effective resolution faces limitations imposed by the size of antibody-fluorophore complexes (Fig. 1B). The finite size of these complexes introduces positional uncertainty (linkage error) by spatially displacing fluorophores from their target epitopes. This limitation is particularly pronounced in the AIS, whose nanoscale architecture contains very closely located structures comparable in size to antibody–fluorophore complexes, resulting in unresolved distances and incomplete labelling due to epitope inaccessibility. Conversely, techniques such as cryo-electron microscopy (cryo-EM) and cryo-tomography (cryo-ET) can achieve sub-nanometre resolution, but without specific labels they provide insufficient target identification within the crowded AIS environment. Platinum-replica EM has shown dense immunogold labelling of proteins within the MPS, such as the C-terminal of 480-kDa AnkG decorating the spectrin mesh and associating with microtubules, yet it has not been able to distinguish the two AnkG molecules predicted to occupy a spectrin tetramer^9–11^. Cryo-EM has yet to fully realize its potential for higher-resolution studies in this context and remains technically challenging.

Expansion microscopy (ExM) has emerged as a powerful super-resolution technique, that overcomes many of these limitations by physically enlarging the specimen^29–32^. Controlled crosslinking and isotropic expansion of the sample allow densely packed features and their individual components to be resolved by improving the effective resolution without the need for changing the hardware of the microscope. Post-expansion labelling protocols such as ultrastructure-expansion microscopy (u-ExM) retain the absolute dimensions of the conventional labelling molecules^33^, but the dimensions of the expanded sample consequently make them relatively smaller, reducing linkage errors and enhancing labelling accuracy. Expansion of the sample prior to labelling also has the effect of de-crowding the target volume, improving epitope accessibility and labelling density, and thereby allowing a more complete mapping of the molecular targets. To date, only a pre-expansion labelling protocol (pro-ExM) has been applied to study the spectrin component of the axonal MPS^34^, leaving u-ExM as an unexploited method in AIS studies.

Here, we report the application of u-ExM (∼4.5x expansion) and *in situ* cryo-ET for studying the nanoscale organisation of the AIS in 3D. U-ExM reveals a consistent ∼80 nm spacing between the C-termini of two adjacent AnkG molecules that reside on a spectrin tetramer interspersed by K_v_1.1 channels. Complementary *in-situ* cryo-ET of flash-cooled neurons confirms that more than two NF-186 molecules are situated at ∼190 nm intervals along the MPS, suggesting a more complex molecular arrangement between NF-186 and AnkG.

Together, these data provide the first direct microscopy evidence for the predicted double-periodicity of AnkG within the canonical ∼190 nm AIS lattice. Our findings refine the macromolecular model of the AIS and its AnkG-associated complex, showing how AnkG pairs can harbour a higher-order scaffold for membrane proteins exemplified by NF-186. Moreover, the study showcases the synergistic power of u-ExM and cryo-ET where ExM provides molecular specificity with nanoscale localization and cryo-ET validates the arrangement in the native, unexpanded state. Together, these approaches push the resolution limits for studying the ultra-dense neuronal ultrastructure and unmask dense features that otherwise remain obscure.

## Materials and Methods

### Neuronal cultures (ExM)

Mixed cultures were prepared from dissociated hippocampal neurons from P0 rat pups. Hippocampi were dissected, homogenized, triturated, and seeded at 30,000-60,000 cells/cm² (40,000-75,000 cells/well) on 13-mm coverlips (#1.5H) coated with poly-D-lysine (PDL; ThermoFisher, cat. No. A3890401). Neurobasal+ medium with B-27+ (ThermoFisher, cat. No. A3653401), Glutamax (ThermoFisher, cat. No. 35050061), and 100 U/mL Penicillin-Streptomycin (ThermoFisher, cat. No. 15140122) was used for seeding and growth. At 2-4 days in vitro (DIV), 10 µM 5-Fluoro-2’-deoxyuridine (FdU; Sigma-Aldrich, cat. No. F0503) was added to inhibit glial proliferation. Half-medium changes were performed 2-3 days later and then twice weekly. Neurons were cultured up to three weeks and fixed in 4% paraformaldehyde (PFA) at 18-21 DIV for expansion microscopy experiments, then stored in PBS at 4 ℃ in the dark.

### Antibodies

The primary antibodies used and the respective dilutions are listed in Supplementary Table S1. Secondary antibodies used were goat anti-rabbit AlexaFluor 568 (1:700, Invitrogen A-11011), goat anti-mouse AlexaFluor 488 (1:700, Invitrogen A-11001) for Kv1.1 and AnkG C-terminal tail (AnkG-C’) labelling, and goat anti-mouse AlexaFluor 568 (1:700, Invitrogen A-11004) and donkey anti-chicken AlexaFluor 488 (1:200, Jackson Immuno 703-545-155) for Kv1.1 and NF-186 labelling in ExM experiments.

### Expansion microscopy with post-expansion labelling

A post-expansion immunolabelling protocol (u-ExM) from Gambarotto et al. (2021) was followed with minor changes. Briefly, 18-22 DIV neurons were fixed with 4% PFA for 10 min, permeabilized in 3% BSA/0.3% Triton-x100 in phosphate-buffered saline (PBS) for 30 min, followed by protein anchoring in 0.7% formaldehyde (FA)/1% acrylamide (AA) in PBS overnight, all at room temperature (RT). For gelation, 35 µL of monomer solution with 0.5% Tetramethylethylenediamine (TEMED) and 0.5% ammonium persulfate (APS) per coverslip was pipetted on parafilm over ice, and the coverslip was inverted onto the droplet, cell side facing down, incubated for 5 min on ice, then 1 h at 37 °C. For mild heat denaturation, gels were incubated in denaturation buffer for 15 min at RT first, and then for 45 min at 95 °C. Gels were expanded in MilliQ water in three rounds, the respective post-expansion diameter was measured for determining expansion factor (ExpF_Gel_), cut into 18-mm pieces (Supplementary Fig. S5), and stored in PBS at 4 °C in the dark until staining.

### Immunofluorescence staining of expanded samples

Staining was performed in Eppendorf tubes in volumes of at least 500 µL. Gels were blocked in 2% BSA in PBS for 2 h at 37 °C, followed by overnight incubation with primary antibodies at 37 °C. Gels were washed three times 10 min each in wash buffer (1% BSA/0.1% Tween20 in PBS) at 37 °C, then incubated with secondary antibodies for 2-3 h at 37 °C. After repeating washes, gels were either stored in PBS at 4 °C or labelled with nuclear stain Hoechst 33342 (1:700, Thermo Scientific 62249) for 20-30 min at RT. All incubation and washing steps were performed with vigorous shaking. Prior to mounting and imaging, gels were expanded in MilliQ water over three rounds of 15 min at RT in the dark. Gels were mounted on PDL-coated 20-mm glass-bottom MatTek dishes, submerged in MilliQ water, and imaged immediately (within 24 h) to minimize fluorescence loss.

### Determination of the expansion factor

ExpF_Gel_ was calculated from the pre- and post-expansion gel dimensions, providing an approximate estimate of the expansion, which is ∼4.5-fold for this protocol. A local expansion factor (ExpF_Kv_) relative to the MPS was determined for individual AIS using the K_v_1.1 periodic organisation in the respective AIS as an internal reference. ExpF_Kv_ was calculated by dividing the mean post-expansion K_v_1.1 inter-peak distance by 190 nm and used to derive pre-expansion AnkG and NF-186 distances (Supplementary Fig. S5B-C). For autocorrelation-based measurements, values from different image planes of the same AIS were treated as replicates and averaged to obtain a single ExpF_Kv_ per AIS. Final ExpF_Kv_ values are reported as mean ± standard error of the mean (s.e.m.), averaged across gels.

### Fluorescence microscopy and image acquisition

Images were acquired using a Zeiss LSM 980 Airyscan 2 inverted confocal laser-scanning microscope with a 63x oil Plan-Apochromat objective (1.4 NA). Excitation was performed with 488 and 543 nm laser lines using GaAsP or Airyscan detectors. Confocal imaging was performed with a pixel size of 85×85 nm² and a pinhole size of 1 AU, bidirectional scanning (speed 9) and 4x line averaging. Airyscan imaging was performed at “super-resolution (SR)” mode with a pixel size of 42.5×42.5 nm², bidirectional scanning (speed 6), and 2x line averaging. Z-stacks were acquired with slice thicknesses of 0.23 µm (confocal) or 0.15 µm (Airyscan). Airyscan images were 3D-processed using auto-filtering in Zeiss ZEN Blue v3.8. Acquisition settings were held constant within experiments and adjusted between experiments to optimize gel-specific signal-to-noise ratio.

### AIS periodicity analysis (ExM)

Autocorrelation analysis was performed using the K2 Napari Wave Breaker plugin v0.1.4^35^ in Napari (doi:10.5281/zenodo.3555620), available at https://github.com/SamKVs/napari-k2-WaveBreaker. Expansion in the Z direction enabled analysis of individual Z stacks with reduced out-of-focus signal, yielding multiple distinct AIS membrane cross-sections. To account for the undulating shape of the AIS membrane in Z, non-overlapping maximum intensity projection (MIP) images were generated in Fiji v2.14.0 (doi:10.1038/nmeth.2019) using ∼10 consecutive planes for Kv1.1 and one to five planes for AnkG or NF-186. Multiple non-overlapping MIPs per AIS were included in the autocorrelation analysis to maximize data usage. A threshold-based mask was applied to the AIS, and grids of 120×80 or 80×120 pixels (5×3.5 µm) were generated (Supplementary Fig. S4A). Grids were analysed across orientations defined by the AIS midline ± 20°. For each grid, an autocorrelation curve was computed, from which the autocorrelation amplitude (height of the first non-zero peak) and the dominant spatial frequency (distance from the origin to the first peak) were extracted automatically (Supplementary Fig. S4B-D). The autocorrelation amplitude reflects the strength of the periodic pattern at the detected frequency.

Analyses were performed using post-expansion measurements. To quantify the characteristic 190-nm MPS periodicity, grids were restricted to a frequency range of 0.7-1.15 µm (∼155-255 nm pre-expansion). Shorter AnkG periodicity was assessed within 0.2-0.6 µm (∼45-130 nm pre-expansion). Frequency values from grids within the same AIS image plane were averaged to obtain mean inter-peak distances for K_v_1.1, AnkG, and NF-186 per AIS. Measurements from different planes of the same AIS were treated as replicates and averaged. Pre-expansion distances were calculated by dividing the mean inter-peak distance per AIS by the AIS-specific ExpF_Kv_, determined from K_v_1.1 reference labelling. Mean pre-expansion distances from all AISs were pooled across independent cultures to derive the final inter-peak distance.

For manual analysis, segmented line ROIs (8-pixel thickness) were manually drawn in Fiji on non-overlapping membrane segments from single-plane AIS images. Intensity profiles were generated, and peaks were detected using the ‘Find Peaks’ tool in the BAR plug-in (v1.5.2; doi:10.5281/zenodo.495245), applying a minimum peak separation of 0.5 µm (∼100 nm pre-expansion). Peak coordinates were exported, and inter-peak distances were calculated. Distances were restricted to 0.7-1.15 µm to define post-expansion K_v_1.1 spacing. Measurements from the same AIS were averaged to obtain a mean inter-peak distance per AIS. For each AIS an ExpF_Kv_ was calculated by dividing the mean inter-peak distance by 190 nm (Supplementary Fig. S5B-C).

For AnkG-C’ labelling, two distance classes were quantified: intra-pair and inter-pair (Fig. 3B-C). Intra-pair distances were defined using the coordinates of two peaks within a single broad peak separated by a valley and flanked by the reference K_v_1.1 signal. Inter-pair distances were defined using the coordinates of two consecutive independent peaks with an intervening K_v_1.1 signal. Individual measurements were divided by the ExpF_Kv_ of the corresponding AIS to obtain pre-expansion distances. Distances from the same AIS were averaged to obtain a mean per AIS, and these values were subsequently averaged across gels.

NF-186 labelling was analysed using the same distance-classification strategy. Inter-pair distances harboured K_v_1.1 signal, which was absent from intra-pair distances. Distances between NF-186 signals observed as close-proximity pairs, oriented either along the proximal-distal AIS axis or more circumferentially (‘closest pairs’; Fig. 4C, E-F), were also measured. These measurements were performed independently of any framing K_v_1.1 signal, although the absence of Kv1.1 within a given pair was required.

### Statistical analysis

For u-ExM experiments, gels from four (K_v_1.1–AnkG) and three (K_v_1.1–NF-186) independent cultures were analysed. Data from all gels within each staining condition were pooled.

All data are presented as mean ± standard deviation (s.d.), except ExpF_Kv_ values averaged across gels (Supplementary Fig. S5B-C), which are reported as mean ± s.e.m. Measurements obtained from the same AIS were averaged to generate AIS-level means used for statistical comparisons where relevant. AIS-level means were pooled across gels to calculate the final reported values.

Statistical analyses were performed using GraphPad Prism v10. Normality was checked using the Shapiro-Wilk test. Comparisons between two groups were performed using two-tailed Wilcoxon matched-pairs signed-rank test (Fig. 2) or two-tailed paired t-test (Figs. 3, 4), as appropriate. Comparisons across three or more groups were performed using the Kruskal-Wallis test (Fig. 5), followed by Dunn’s multiple comparisons test with Bonferroni correction. Statistical significance is shown as: ****P < 0.0001.

**Figure 2.**
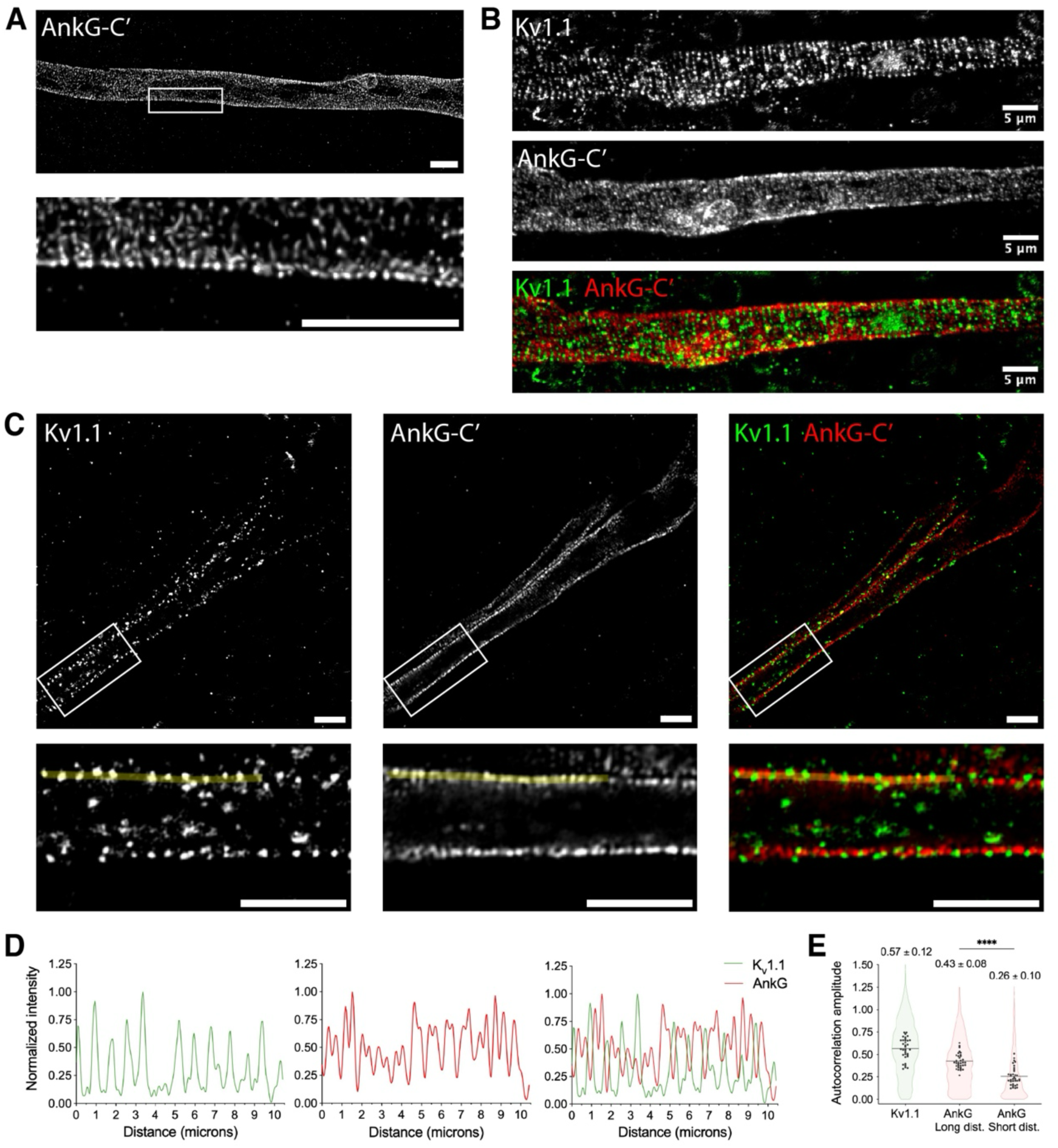
Paired AnkG-C’ labelling interspersed with K_v_1.1 along the AIS. **A,** Post-expansion labelling of the AnkG-C’ reveals paired AnkG signals in the expanded AIS. Maximum intensity projection (MIP) of an Airyscan image. **B-C**, MIP confocal (**B**) and single-plane Airyscan (**C**) images showing interspersed K_v_1.1 (green) and AnkG-C′ (red) labelling along the membrane of an expanded AIS. **A-C**, Scale bars, 5 µm (post-expansion). **D**, Intensity profiles of fluorescence signals along the yellow lines in **C** reveal distinct patterns for the two channels. K_v_1.1 profile (green) shows single peaks with ∼190 nm spacing, whereas AnkG-C′ profile (red) displays two partially resolved peaks within each broader peak, intercalated by K_v_1.1 signal. Distances shown correspond to post-expansion measurements. **E**, Autocorrelation amplitudes (AC) of K_v_1.1 and AnkG-C′ labelling as indicators of periodicity strength. Truncated violin plots represent grid-level AC values, and black points indicate AIS-level mean AC values used for statistical analysis. Strong periodicity is detected for both K_v_1.1 and AnkG-C′ when analysed within a post-expansion frequency range of 0.7-1.15 µm. AnkG-C′ periodicity is weaker when the analysis is restricted to a frequency range of 0.2-0.6 µm. Number of grids plotted: N = 1,587 (K_v_1.1), N = 1,562 (AnkG Long distances), and N = 1,441 (AnkG Short distances). Data are shown as mean ± s.d.; n = 35 AIS from 4 independent cultures. Two-tailed Wilcoxon matched-pairs signed-rank test (AnkG Long vs Short distances), **** P < 0.0001.

### Neuronal culture (EM)

Cryo-EM grids were prepped for cell culture by glow discharging with the PELCO easiGlow Glow Discharge Cleaning System at the cryo-EM facility EMBION. Grids were dipped in 70% ethanol for further sterilization and placed in 35-mm Mattek dishes on glass surface. Grids and glass coverslip of the Mattek dish were coated with 40 μg/mL poly-d-lysine (ThermoFisher, cat. No. A3890401) for 1h at RT and then washed in sterile water 3 times. For the last wash the water was replaced with neuronal plating medium and incubated at 37℃ and 5% CO_2_ overnight. Primary rat hippocampus neurons were cultured on cryo-EM grids with a density of approx. 50.000 cells/grid. Dissociated cells in a high concentration were added to the pre-heated growth medium on the grids and were left to attach for 1-3 h before supplementing with 2 mL medium. Two to four days after seeding anti-mitotic drug 5-Fluoro-2’-deoxyuridine (FdU; Sigma-Aldrich, cat. No. F0503) was added to the medium at 10 μM final concentration to suppress glial cell growth. Half of the medium was changed after two-three days and afterwards the cultures were maintained by changing half of the medium with fresh growth medium twice a week. Cultures were kept until 14-16 DIV before live-labelling and plunge freezing and then stored in liquid nitrogen until cryo-imaging.

### Live labelling

Neuronal growth medium was removed from the cells, and the cells were incubated with primary antibody solution in a 1:100 v/v dilution in growth medium for 30 min at 37℃ and 5% CO_2_. The cells were washed three times in prewarmed medium. Secondary antibody (anti-mouse IgG gold antibody #G7652 Sigma Aldrich) was diluted in neuronal medium 1:20 v/v and 200 µL was added to the cells and incubated for 15 min at 37℃ and 5% CO_2_. The cells were washed 3 times in prewarmed medium. Hereafter, the grids were taken directly to the plunge freezer to limit dissociation and/or uptake of the antibodies.

### Cryo-EM grid preparation

Plunge freezing of cell cultures on EM grids were performed on a Leica GP2 plunge freezer at EMBION. The living culture was transported in a CELLBOX2.0™, and grids were plunge frozen within 5 min from being taken out of the incubator. Grids were picked up directly from culture dishes, and 3 µL prewarmed PBS was added to the front before back-blotting for 8 sec. The Leica GP2 was operating at 25℃ and 95% relative humidity while plunging. Grids were plunged into liquid ethane kept at -184℃. Grids were then transferred to a grid storage box and stored in liquid nitrogen until subjected to cryo-ET experiments.

### Cryo-EM data acquisition

Micrographs and tilt series were acquired on a Titan Krios G3 (Thermo Fischer) operating at 300 kV using the SerialEM software pakage [60]. Low-magnification montages were used for navigation and search purposes, and areas of interest was chosen based on the localization of neurites, cell density, ice thickness and guided by gold labelling. Micrographs and tilt-series data were collected at regions of interest at 26,000x magnification, with a resulting pixel size of 3.4 Å. Low-dose images and tilt series, with defocus values of -8 µm, were recorded with the Gatan K2 Summit camera using an energy filter with a slit width of 30 eV. Movies of 11 frames were collected at an angular tilt range from -60° to +60° using a dose-symmetric collection scheme with an increase of 2° per image starting at 0°. The total electron dose for a complete tilt series was kept below 120 e^-^/Å^2^.

### Gold labelling analysis

In attempts to analyze the clustering of gold particles in the micrographs, no existing software was found to solve the problem. Thus, a python-script with a graphic user interface (GUI) integrating image-alignment and clustering algorithms was developed and is presented here. The created program recognizes original and binary filtered images with the same names and overlay them after performing clustering by DBSCAN algorithm on the binary image. The GUI shows the overlay image and labels the clusters with the number of particles in each cluster as shown in figure 4. Left clicking on the clusters will output the distance between the selected clusters in pixels. The code and the user manual are available at: https://github.com/jkdannerso/Cluster-analysis-cryoEM-2D.git.

### Cryo-ET data processing

Tilt series movies were motion corrected using MotionCor2^36^. The motion corrected tilt series were aligned and tomograms reconstructed using IMOD^37^ and the semiautomated Etomo software^38^. Gold-labelling was picked by using IMOD findbeads3d, outputting a model file and 3D coordinates of the single gold particles in the tomograms. A 3D clustering analysis was performed, running the DBSCAN algorithm for clustering as implemented in the sklearn.cluster package in Python3. The maximum allowed distance between points within a cluster (eps) was set at 20 pixels (54.4 nm) and minimal cluster size was set at two gold particles. This value was determined based on visual inspection of the clustering showing distinct clusters with no overlap. The cluster sizes and widths were extracted by a home-build python-script (available at https://github.com/jkdannerso/Cluster-analysis-3D.git), clusters were categorized by size (number of gold particles, n), and the cluster width (maximum distance from point to point in the cluster). A total of 219 clusters of sizes between 2-16 particles were detected and analysed, and 14 clusters was removed from the dataset as outliers based on their size using a modified Z-score > 3.5.

## Results

### Post-expansion labelling of the AIS enables nanoscale visualisation of the MPS

The ∼4.5-fold lateral (∼90-fold volumetric) expansion of the samples achieved with u-ExM protocol results in a comparable improvement in the effective resolution of our imaging setup (Airyscan confocal microscope) using 488 and 568 nm wavelengths. The theoretical lateral resolution of our microscope with an oil objective lens (NA=1.4) for 568 nm wavelength is 247 nm according to the Rayleigh criterion^39^ for confocal mode and reported as 120 nm for Airyscan mode using 488 nm wavelength^40^. Using an expansion factor of ∼4.5, the nominal spatial resolution becomes ∼55 and ∼27 nm (247 nm or 120 nm divided by 4.5) for confocal and Airyscan modes, respectively, which are sufficient to visualize the 190-nm spatial period of the MPS without the need for any additional optical components for super-resolution microscopy (Fig. 2 and Supplementary Fig. S2). Using an antibody targeted at the C-terminal tail of AnkG (AnkG-C’) for post-expansion labelling, we could clearly observe a two-layered pattern of AnkG that was conserved along the AIS in the expanded sample (Fig. 2A), in addition to the well-defined uniform ∼190-nm spatial period of the SBD of AnkG^9^.

### AnkG exhibits a strict two-molecule organisation per MPS repeat

We speculated that the pairs of AnkG signal that are clearly visible in both confocal (Supplementary Fig. S2) and Airyscan images (Fig. 2A) could be the C-terminal tails of AnkG pairs localized on the same spectrin tetramer. To check this, we looked for an AIS component whose labelling would intercalate the AnkG pairs as a whole and occupy the space between AnkG pairs localized on adjacent spectrin tetramers. Although actin rings are known to intercalate the 190-nm spatial period of AnkG in the AIS, phalloidin-based actin labelling has been reported to be incompatible with standard ExM protocols^41^. We therefore selected K_v_1.1 as an alternative marker. K_v_1.1 is known to associate with K_v_1.2^42,43^, which is periodically organised along the AIS and colocalizes with actin rings^44^. Using an antibody against K_v_1.1, we confirmed that K_v_1.1 colocalizes with K_v_1.2 and together they intercalate NF-186 in pre-labelled and expanded AIS (Supplementary Fig. S3). Proceeding with this antibody, the post-expansion labelling of K_v_1.1 and AnkG-C’ revealed a punctate K_v_1.1 pattern emerging from the spaces between AnkG pairs (Fig. 2B-D) consistent with our prediction. These findings confirm that the paired AnkG signal arises from the C-terminal tails of AnkG molecules associated with a spectrin tetramer within each MPS repeat.

Estimating native distances from measurements made on expanded samples requires accurate determination of the expansion factor (ExpF). Conventionally, ExpF_Gel_ is calculated as the ratio of post-expansion to pre-expansion gel dimension. However, local inhomogeneities, arising for example from variations in protein content, can introduce distortions that limit the reliability of this global estimate^34^. To obtain a local and more accurate ExpF, we used the ∼190-nm spatial period of the MPS as an internal reference. The singular fluorescence pattern of K_v_1.1 labelling within a MPS repeat enabled a more robust estimation of the post-expansion MPS periodicity compared to the AnkG-C’ labelling, where the paired signal complicates distance measurements. Accordingly, K_v_1.1 labelling was used as the reference for ExpF determination in subsequent experiments (Supplementary Fig. S5B-C).

To quantify K_v_1.1 and AnkG-C’ labelling in the MPS of expanded samples, we performed an autocorrelation analysis^45,46^ (Fig. 2E, Supplementary Fig. S4). The mean autocorrelation amplitude (AC) for the reference K_v_1.1 labelling was 0.57 ± 0.12 at a post-expansion mean inter-peak distance of 894 ± 40 nm (Fig. 2E). For each AIS, a corresponding ExpF_Kv_ was calculated by dividing the mean post-expansion inter-peak distance of the K_v_1.1 labelling by 190 nm. ExpF_Kv_ exhibited variability across AIS and gels (Supplementary Fig. S5B). To account for this variability, ExpF_Kv_ determined for each AIS was used to calculate the corresponding pre-expansion distances of AnkG and NF-186 labelling. Using this approach and a post-expansion frequency range of 0.7-1.15 µm, a mean inter-peak distance of 188 ± 7 nm (pre-expansion) with a mean AC of 0.43 ± 0.08 was obtained for the larger patterns of AnkG-C’ labelling (Fig. 2E), consistent with the ∼190-nm spatial period of the AIS. In an attempt to estimate the shorter distances suggested by the AnkG-C’ intensity profiles (Fig. 2D), we restricted the autocorrelation analysis to a post-expansion frequency range of 0.2-0.6 µm. This analysis resulted in a mean inter-peak distance of 103 ± 7 nm (pre-expansion) with a significantly lower mean AC (0.26 ± 0.10; two-sided Wilcoxon matched-pairs signed-rank test, n = 35, P < 0.0001; Fig. 2E). This value may be a good approximation of the shorter distances between AnkG-C’ labelling, corresponding to approximately half of the ∼190-nm period of the MPS. However, by visual inspection, we observed that the paired AnkG-C’ signals exhibited non-uniform spacing, with shorter distances observed within pairs than between pairs (Fig. 2A-D). Such heterogeneity cannot be resolved by autocorrelation analysis, which inherently assumes a single dominant periodic pattern.

### AnkG C-termini organise pairwise in the MPS

To dissect this pattern and distinguish the two distances, we performed manual calculations on the same set of AISs (Fig. 3). Using K_v_1.1 signal as the reference, the distances between the AnkG-C’s were classified as intra- or inter-pair which yielded the respective mean distances of 80 ± 4 nm (intra-pair) and 104 ± 4 nm (inter-pair; Fig. 3D). Remarkably, these two subpopulations consist of partially overlapping range of distances (50-110 nm intra-pair and 69-139 nm inter-pair distances; mean ± 2x s.d. of individual measurements). The overlap between these subpopulations makes their separation difficult without the visual inspection in the presence of another component such as K_v_1.1. Despite the variability of AnkG-C’ positioning in the AIS, the distance within an AnkG pair on a spectrin tetramer is significantly shorter on average than the distance between two pairs (two-tailed paired t-test, t (32) = 19.47, P < 0.0001; Fig. 3D). This indicates a two-fold periodic localization of AnkG-C’s and extends our current understanding of its organisation in the AIS.

**Figure 3.**
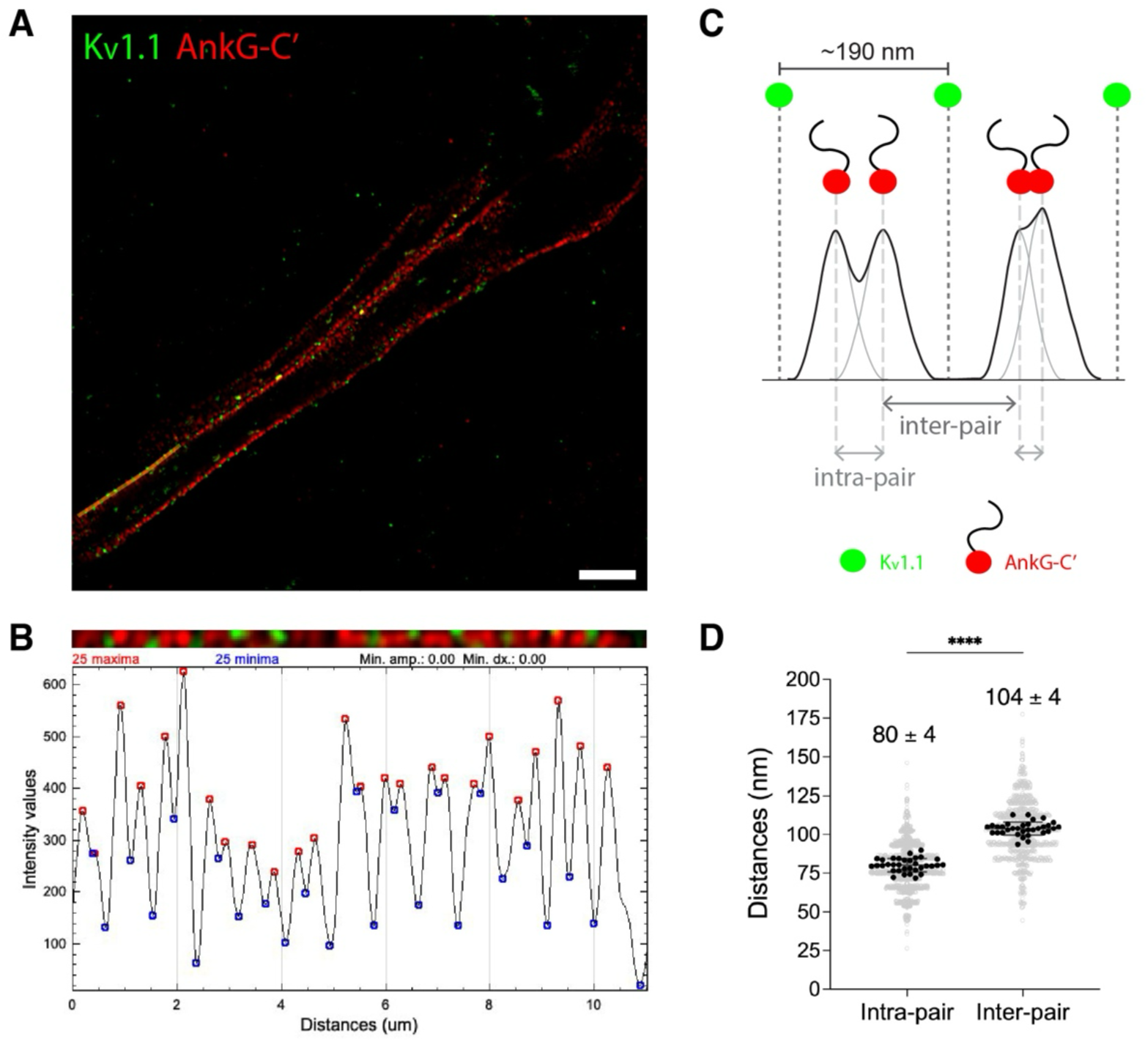
Manual analysis of paired AnkG-C’ labelling. **A,** Single-plane Airyscan image of an expanded AIS labelled for K_v_1.1 (green) and AnkG-C′ (red). Scale bar, 5 µm (post-expansion). **B**, Close-up of the yellow line ROI in **A** and the corresponding AnkG-C′ fluorescence intensity profile. Peaks and valleys (red and blue circles, respectively) were marked using the ‘findpeaks’ tool of the BAR plug-in in Fiji. Based on visual inspection of the labelling along the line ROI, distances between peaks were measured and classified as intra- or inter-pair distances. **C**, Schematic representation of intra- and inter-pair peak distances based on overlapping PSFs of AnkG-C’ labelling. Left, AnkG pairs separated by distances above the effective resolution produce two resolved peaks (PSFs) with a valley between them. Right, AnkG pairs separated by distances below the effective resolution produce overlapping peaks (PSFs) without a clear valley which obscures the underlying signal. Intra-pair distances correspond to the distance between two peaks without an intervening K_v_1.1 signal, whereas inter-pair distances correspond to peaks separated by a K_v_1.1 signal. **D**, Individual intra- and inter-pair distance measurements (pre-expansion values) obtained from manual analysis are shown in grey. Black points represent AIS-level mean values used for statistical analysis. Data are shown as mean ± s.d.; n = 33 AIS from 4 independent cultures. Two-tailed paired t-test, t(32) = 19.47, **** P < 0.0001.

### Multiple NF-186 molecules organise along AnkG pairs in the MPS

To show if NF-186 exhibited a two-fold organisation in the AIS by associating with the two AnkG molecules within each ∼190-nm spatial period, we did post-expansion labelling with antibodies against K_v_1.1 and NF-186 in the AIS (Fig. 4). Autocorrelation analysis of NF-186 labelling using a post-expansion frequency range of 0.7-1.15 µm confirmed its periodic organisation in the expanded MPS with a mean inter-peak distance of 184 ± 9 nm (pre-expansion; AC: 0.45 ± 0.051). By contrast, using a post-expansion frequency range of 0.2-0.6 µm resulted in a mean inter-peak distance of 101 ± 4 nm (pre-expansion) with a significantly lower mean AC of 0.33 ± 0.051, (two-tailed paired t-test, t (13) = 10.5, P < 0.0001; Fig. 4B). Following the same approach as for AnkG-C’ labelling, we continued with manual calculations (Fig. 4D-E), which confirmed the paired organisation of NF-186 with a mean intra-pair distance of 77 ± 6.0 nm, significantly shorter than the distance between the pairs (inter-pair; two-tailed paired t-test, t (13) = 9.846, P < 0.0001; Fig. 4E). This is well in line with the AnkG-C’ intra-pair distance of ∼80 nm. However, the paired organisation of NF-186 deviated more compared to the paired AnkG signal. On occasions, as few as one or as many as four NF-186 puncta were observed within a ∼190-nm spatial period framed by the Kv1.1 signal (Fig. 4A, F). This was corroborated by the observation of close-proximity NF-186 pairs (closest pairs) that were spaced apart 52 ± 2.6 nm, significantly shorter than intra-pair distances (two-tailed paired t-test, t (13) = 14.41, P < 0.0001; Fig. 4E), representing the smallest distance between NF-186 pairs that can be observed with the post-expansion labelling ExM used in this study. These NF-186 pairs were not strictly aligned with the proximal-distal axis of the AIS and were oriented circumferentially as well (Fig. 4C, yellow vs cyan arrowheads). Occasional observation of these pairs in clusters of three to four puncta within a ∼190-nm spatial period framed by K_v_1.1 labelling (Fig. 4A, marked with an asterisk) hints at a multi-molecule organisation of NF-186 with AnkG in the AIS. In accordance, the observation of up to four NF-186 puncta within a ∼190-nm spatial period (Fig. 4F) can be explained by two pairs of NF-186 associated with the two adjacent AnkG molecules on a spectrin tetramer. This complex molecular organisation of NF-186, based on two distinct NF-186 binding sites on the MBD of AnkG as previously shown^47^, can explain why its organisation shows high variability and appears periodic at varying extents as illustrated in Supplementary Fig. S6 and previously in other super-resolution microscopy studies^9^.

**Figure 4.**
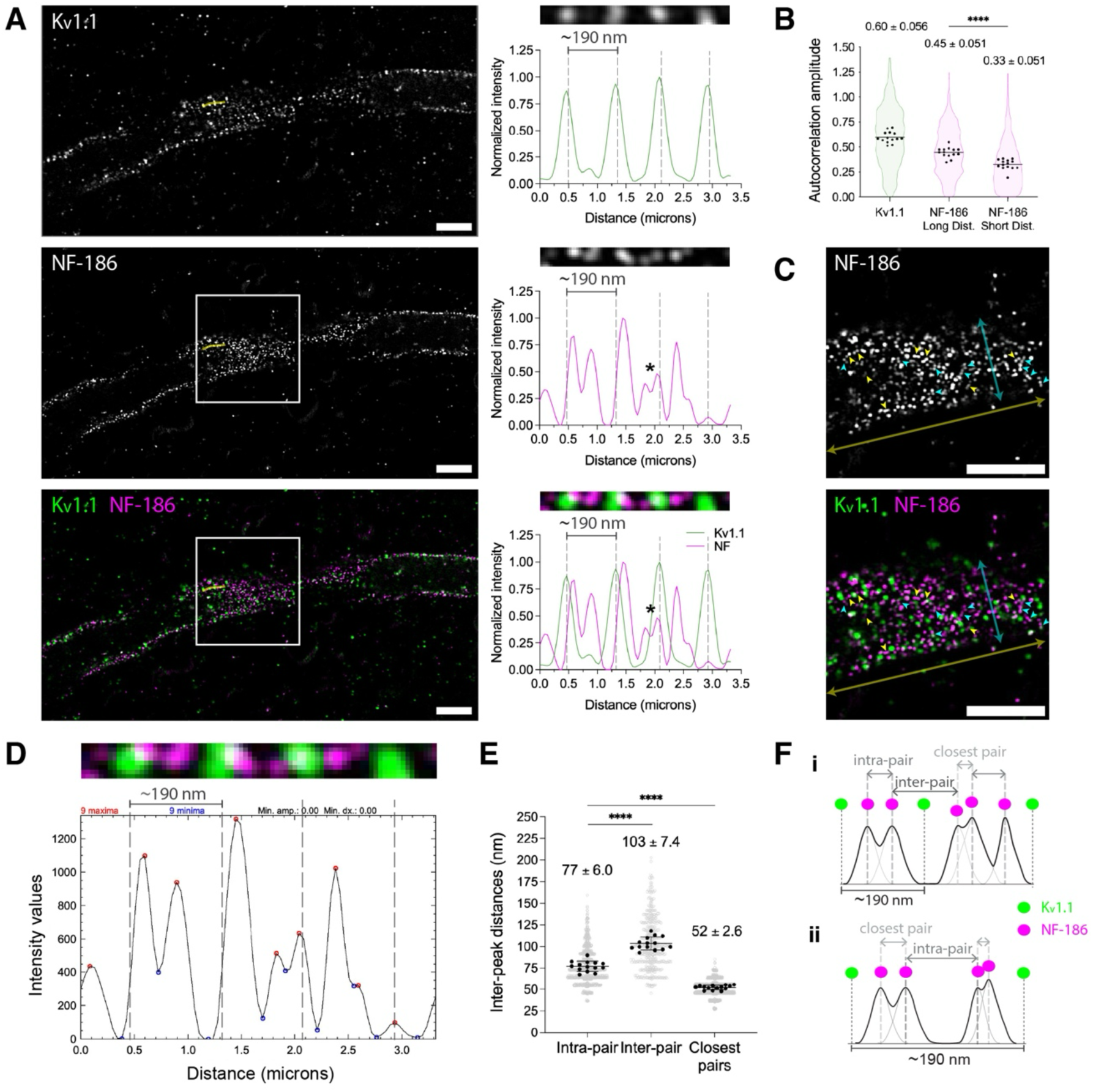
Manual analysis of NF-186 labelling in the AIS. **A,** Single-plane Airyscan images of an expanded AIS showing interspersed K_v_1.1 (green) and NF-186 (magenta) labelling along the membrane (left) and corresponding fluorescence intensity profiles along the yellow line ROI (right). K_v_1.1 profile shows single peaks, whereas NF-186 profile displays 1-4 peaks within each ∼190-nm spatial period defined by K_v_1.1 (dashed lines). Asterisk-marked peaks may represent two NF-186 molecules associated with one AnkG. Distances are post-expansion. **B**, Autocorrelation analysis of K_v_1.1 and NF-186 signals. Truncated violin plots show autocorrelation amplitudes analysed for longer (0.7-1.15 µm) and shorter (0.2-0.6 µm) post-expansion frequency ranges. N = 701 grids (K_v_1.1), N = 614 grids (NF-186 Long distances) and N = 709 grids (NF-186 Short distances). Data are mean ± s.d.; n = 14 AIS from 3 independent cultures. Two-tailed paired t-test, NF-186 Long vs Short distances, t(13) = 10.5, **** P < 0.0001. **C**, Close-up on regions from **A** (white boxes) showing NF-186 closest pairs (arrowheads; the smallest NF-186 distances resolved in u-ExM) in various orientations relative to the AIS axis. Pairs aligned with the proximal-distal AIS axis (yellow arrow) are marked by yellow arrowheads, more circumferentially (cyan arrow) oriented pairs are indicated by cyan arrowheads. Scale bars, 5 µm (post-expansion; **A, C**). **D-E**, Manual measurements of NF-186 intra-pair, inter-pair and closest-pair distances. **D**, Peaks (red) and valleys (blue) were detected using the ‘findpeaks’ tool of the BAR plug-in (Fiji). Vertical dashed lines indicate the K_v_1.1 defined pre-expansion ∼190-nm spatial period along the AIS. Intensity profiles show post-expansion distances. **E**, Individual measurements (grey; pre-expansion) and AIS-level mean values (black) used for statistics. Data are mean ± s.d.; n = 14 AIS from 3 independent cultures. Two-tailed paired t-test, intra-pair vs inter-pair, t(13) = 9.846, **** P < 0.0001; intra-pair vs closest pairs, t(13) = 14.41, **** P < 0.0001). **F**, Schematic of intra-pair, inter-pair and closest-pair distance measurements based on overlapping PSFs of NF-186 labelling. Examples show ∼190-nm K_v_1.1-defined periods containing two, three (F-i) or four (F-ii) NF-186 peaks associated with two AnkG molecules. Configurations with four peaks may also occur in F-i but remain unresolved due to the effective resolution limit.

### Cryo-EM confirms complex organisation of NF-186 in the AIS

To confirm the observation of a multi-molecule organisation of NF-186, cryo-electron microscopy experiments were performed. Extracellular NF-186 labelling of live neurons worked well for AIS localization in cryo-electron microscopy studies (Fig. 5A-F). The immunogold labelling appeared in clusters of gold particles decorating the plasma membrane in a periodic pattern when analysed in 2D-projection images (Fig. 5B-C). Analysis of the inter-centroid distances of immunogold labelling of intact neurons in cryo-EM resulted in a mean distance of 199 ± 46.5 nm (Fig. 5C) fitting well with the earlier reported spatial period of NF-186 in the AIS^9^, thus showing that the immunogold successfully labels the NF-186 at the plasma membrane of the AIS. Furthermore, we performed cryo-ET experiments with the same samples and found that the 10-nm gold clusters distribute in different Z heights along the plasma membrane (Fig. 5E) and vary in number of particles in a cluster (“size”) and maximum distance across the cluster (“width”). From 19 tomograms, 219 clusters of sizes of 2-16 particles were found, 90,9% of which form clusters of 2-8 particles. The width of the clusters increases by cluster size, and three distinct populations were analysed as 2, 3-4, and 5-8 gold particles per cluster, resulting in cluster widths of 29.3 ± 10.8 nm, 51.8 ± 16.0, and 78.4 ± 19.6 nm, respectively (Kruskal-Wallis test with Dunn’s post-hoc comparisons, H(2) = 141.8, All comparisons P < 0.0001; Fig. 5F), comparing well with the distances for NF-186 observed by ExM.

**Figure 5.**
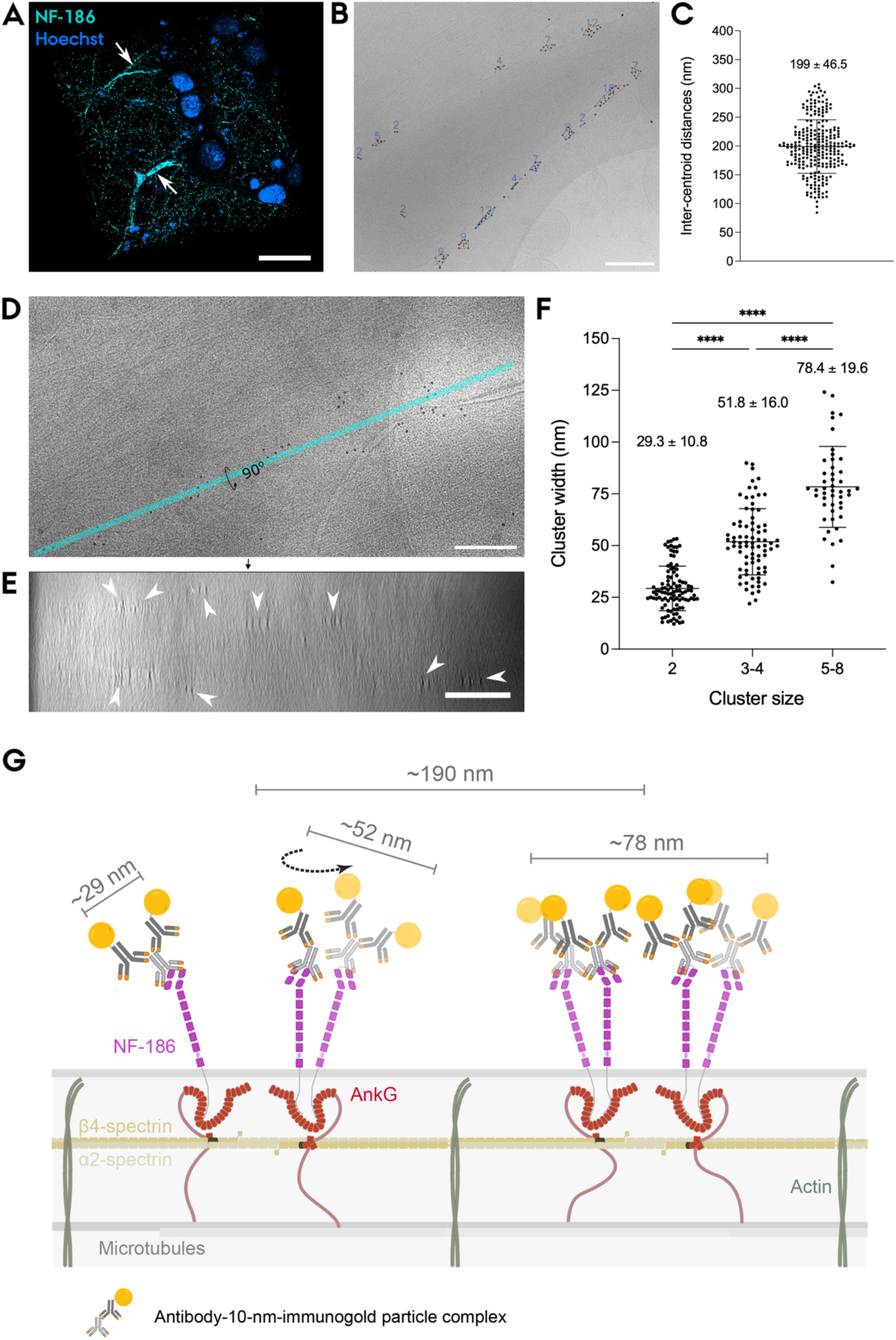
NF-186 clustering and distance analysis from cryo-EM. **A,** Live fluorescence labelling of NF-186 in neurons cultured on EM grids. Arrows point to NF-186-labelled AISs (cyan). Scale bar, 20 µm. **B,** Cryo-EM projection image with DBSCAN clustering of gold particles overlaid. Numbers indicate the count of particles per cluster; red dots mark cluster centroids.**C,** Inter-centroid cluster distances with a mean of 199 ± 46.5 nm (mean ± s.d.). N = 254 distances from n = 45 projection images across 4 independent cultures. **D,** 0° tilt image from a tilt series showing the gold labelled NF-186 decorating the plasma membrane of the AIS. The cyan bar indicates the x-axis shown in the 90° tilted view in **e**. **e,** A 200-nm-thick YZ slice through the reconstructed tomogram showing gold labelling along the plasma membrane and clusters of gold particles (arrowheads) distributed at different Z heights within the cellular volume. Scale bars, 200 nm (**B, D, E**). **F,** Cluster size analysis of the gold particles in 3D. Clusters grouped to be containing 2, 3-4, or 5-8 particles (“size”) show maximum distance between two particles in a cluster (“width”) of 29.3 ± 10.8 nm, 51.8 ± 16.0 nm and 78.4 ± 19.6 nm (mean ± s.d., N=219 clusters, n=19 tomograms from 3 independent cultures.). Kruskal-Wallis test with Dunn’s multiple-comparisons test, H(2) = 141.8, All comparisons **** P < 0.0001. **G,** Schematic illustrating antibody binding and gold cluster formation. A primary antibody (light grey) bound to a NF-186 molecule can recruit one to two secondary antibodies (dark grey) conjugated to a 10-nm gold particle. One NF-186 can therefore produce clusters of up to two gold particles; two NF-186 molecules up to four particles; and 3–4 NF-186 molecules associated with two AnkG molecules up to 5–8 particles. The arrow indicates rotation in the plane of the two NF-186 molecules bound to one AnkG, resulting in varying orientation of the detected clusters in relation to the plane of the plasma membrane.

As the labelling for cryo-EM was performed with a monoclonal primary antibody and a polyclonal secondary antibody, we cannot deduce the actual number of NF-186 molecules represented by these clusters of gold particles. However, the labelling shows a clear tendency to arrange in pairs of gold particles with a very narrow distance. This is also clear from a nearest neighbour analysis of the entire dataset of 1636 Nearest-neighbour-pairs, showing a peak value at 23.8 ± 6 nm with 75% of the total particles having a nearest neighbour within 35 nm distance (Supplementary Fig. S8). Assuming that one or two secondary antibodies can bind one primary antibody, the distance of 23.8 nm fits well with the assumed distance between two secondary antibodies conjugated with 10-nm gold. Following this model, the clusters of size n=2 might represent single NF-186 molecules, clusters of size n=3-4 might represent two molecules of NF-186, and clusters of size n=5-8 might represent three to four molecules of NF-186 (Fig. 5G). Matching this to the ExM data, it shows that NF-186 is periodically arranged along the AnkG-associated complex in the AIS, as described in previous fluorescence microscopy studies^17,47^, while also showing that more than two NF-186 molecules occupy ∼190 nm intervals. This indicates multi-molecule organisation of NF-186 along AnkG in the AIS.

These inter-related distances lead us to a more detailed architecture of the AIS. We incorporate ∼80 nm distance between the C-terminal tails of an AnkG pair located on a spectrin tetramer into the known model of AnkG organization in the AIS with a ∼190-nm spatial period. Furthermore, the close-proximity NF-186 pairs (∼52 nm) and the ∼77 nm spacing between the adjacent pairs framed by Kv1.1 are incorporated into this model and define how one to four NF-186 molecules can associate with an AnkG pair within a ∼190-nm spatial period of the AIS. Together, these measurements establish a refined nanoscale framework for the organisation of AnkG and its associated membrane proteins in the AIS (Fig. 6).

**Figure 6.**
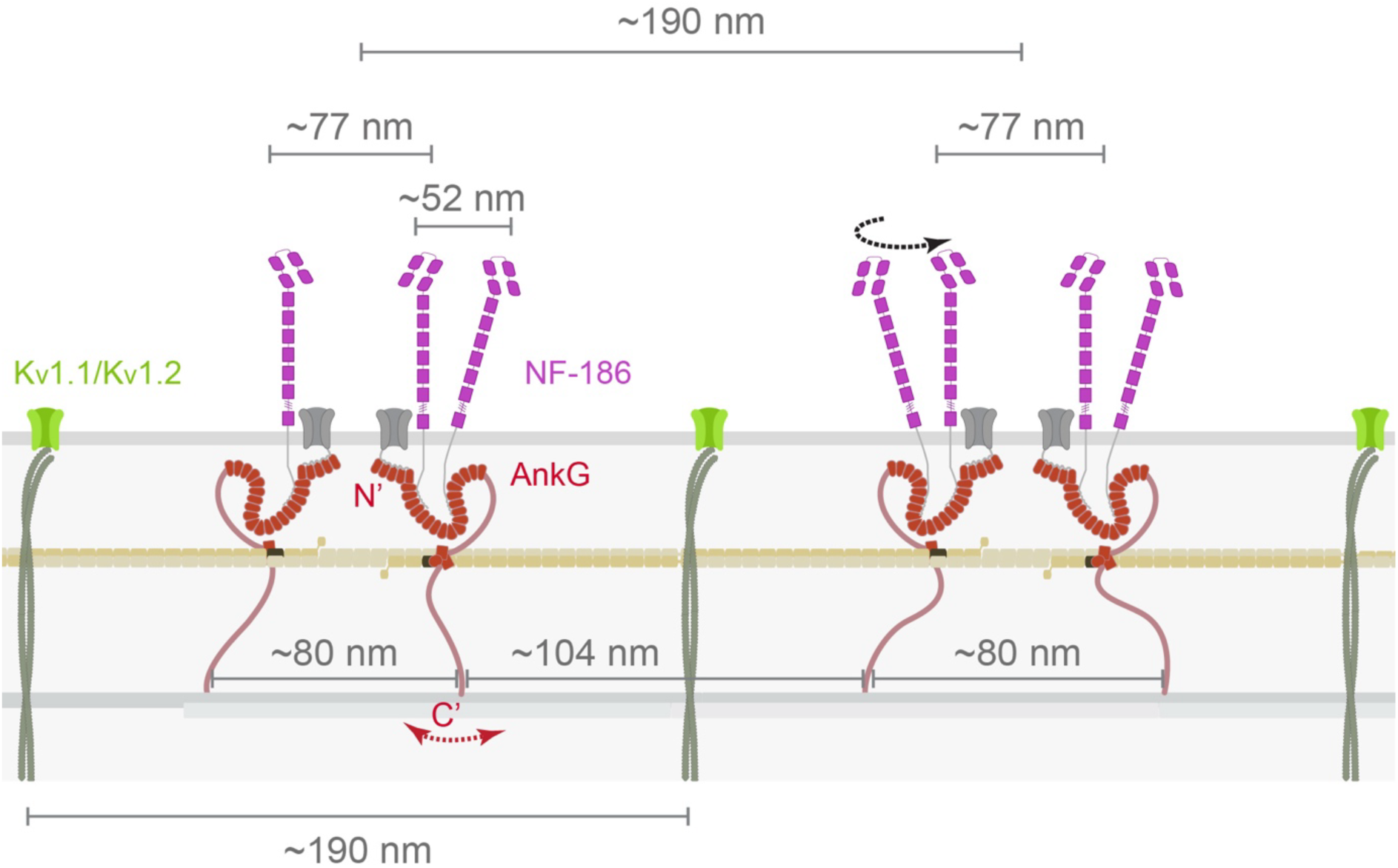
Comprehensive nanoscale model of the AIS incorporating newly resolved distances related to the AnkG-associated complex. ∼190-nm spatial period is observed for K_v_1.1/K_v_1.2 (green) and the AnkG-associated complex (centre-to-centre). Newly resolved AnkG (red) intra- and inter-pair distances are shown as ∼80 nm and ∼104 nm, respectively, measured between C-terminal tails (C’) bound to the microtubules. Flexibility in C-terminal tail positioning is indicated by the red arrow. Through association with AnkG, NF-186 (magenta) adopts a paired organisation, indicated by ∼77 nm spacing between molecules positioned on adjacent AnkG molecules within ∼190-nm spatial period. In addition, more than one NF-186 molecule can associate with a single AnkG molecule, yielding more than two NF-186 molecules per ∼190-nm spatial period. The closest NF-186 pairs (∼52 nm spacing) likely represent molecules associated with the same AnkG molecule. The black arrow indicates multiple orientations (proximal-distal axis or circumferential) observed for these pairs on the plasma membrane. Distances are not drawn to scale.

## Discussion

The 190-nm periodicity of the MPS has been well studied for the main repeating complexes in the AIS such as AnkG, β-IV-spectrin and actin rings. However, restricted by resolution and incomplete labelling, further exploration of finer organisation of the AIS components has remained limited. Overcoming these technical limitations with the use of ExM and cryo-ET, our study provides new insights into the macromolecular organisation of several key components of the AIS.

We show that u-ExM is a well-suited super-resolution fluorescence microscopy method for studying the MPS by validating the periodic arrangement of K_v_1.1, AnkG and NF-186 in the expanded samples. We used a post-expansion labelling protocol^30,33^ with ∼4.6-fold actual expansion which made it possible to visualize the MPS by Airyscan microscopy or even by a conventional confocal microscope. Previously, a pre-expansion labelling protocol was used to visualize axonal MPS for reliable quantitative analysis^34^. Although ExM with both pre- and post-expansion labelling protocols rely on the physical expansion of a sample to achieve super-resolution precision and are sufficient to visualize the MPS, post-expansion labelling provides several additional advantages. It results in higher labelling efficiency as expansion creates a less crowded environment for labelling molecules to penetrate more easily and possibly revealing epitopes that had been previously inaccessible to labelling molecules. After expansion, the size of the labelling molecules becomes effectively smaller compared to the structures to be labelled, decreasing the linkage error resulting from the additional size of the labelling molecules. These represent in fact the serious limitations of current optical super-resolution techniques. The effective improvement in spatial resolution brought on by ExM also extends equally into the Z axis, which otherwise suffers from lower resolution compared to the X and Y directions in optical imaging.

As discussed by Martinez et al. (2020), the use of ExM for quantitative measurements requires accurate determination of the ExpF of the setup. Expansion of a gel depends on several factors such as fixation, denaturation, and the formula of anchoring and gelation solutions etc. Even though each protocol provides an approximate ExpF, the extent of expansion in different areas of the gel and of individual cellular areas and structures can be quite variable. Martinez et al. (2020) showed that gels containing embedded cells expand more compared to the gel itself. We, too, observed slightly higher ExpF (within a range of 4.49-4.83x; Supplementary Fig. S5) than the original 4.5-fold premised by the u-ExM protocol used^33^ and the measured ExpF_Gel_ in this study. Furthermore, we observed slight variations in expansion in the AIS of different neurons within the same gel (Supplementary Fig. S5). We therefore reasoned that the expansion in the ∼190-nm spatial period of an established AIS component could be used to provide a ‘locally accurate’ ExpF for each AIS.

We showed that K_v_1.1 periodically organises in the AIS and co-localizes with K_v_1.2 every ∼190-nm as shown previously^44^, and together they intercalate NF-186 (Supplementary Fig. S3). Thus, we used K_v_1.1 labelling as an internal yardstick for the expansion throughout this study. We calculated ExpF_Kv_ for each AIS using the post-expansion inter-peak distances of K_v_1.1 labelling and used this for determining the pre-expansion distances of the AnkG C-terminal and NF-186 organisation in the respective AIS. While this approach may provide a more appropriate ExpF, it is not without caveats given the possibility of differential expansion of AIS components (intracellular, extracellular, complexes etc.).

Our results from ExM fluorescence data using autocorrelation analysis confirmed the existing periodic localization of AnkG with ∼188 nm in the MPS of expanded neurons (Fig. 1), while manual calculations revealed the presence of a second layer of periodic pattern with shorter distances. Exploiting the advantages of u-ExM and using K_v_1.1 as a reference, we resolved the C-terminal tails of an AnkG pair closely located on a spectrin tetramer and separated these from the C-terminal tails of the neighbouring AnkG pairs (Fig. 2), confirming that exactly two AnkG molecules are situated within ∼190-nm spatial period of the AIS. We distinguished the two sub-populations of distances that define the intra- and inter-pair spacing of two AnkGs bound to spectrin tetramers. We observe that AnkG intra-pair distance (two AnkG C-termini on the same tetramer) range between 50-110 nm with a ∼80-nm mean, and the inter-pair distance (AnkG C-termini from molecules on adjacent tetramers) range between 69-139 nm with a ∼104-nm mean. These sub-populations partially overlap possibly due to the flexibility of the AnkG-C’, which occasionally appears as fluctuating intra-pair distances towards the upper end of the 50-110 nm range. Our data provide an explanation for the observed loss of periodicity for the C-terminal labelling observed in earlier super-resolution microscopy studies^9^. Rather than being highly flexible, as earlier described, the C-terminal appears relatively constrained in relation to the canonical AIS periodicity, potentially stabilised by its interaction with the microtubules through end-binding protein 1 and 3^48^. This may be important for further structural studies to identify a possible function of periodically ordered binding of the AnkG-C’ to microtubular fascicles of the AIS.

Elaborating on the AnkG-associated complex, we investigated the organisation of NF-186 by post-expansion labelling together with K_v_1.1. Autocorrelation analysis of NF-186 on expanded AIS confirmed that NF-186 is organised with a spatial period of ∼184 nm as previously shown. Strikingly, further analysis revealed that NF-186 does not only follow AnkG pairs with ∼77 nm intra-pair spacing (Fig. 4E), but also up to two NF-186 molecules (closest pairs) can associate with one AnkG which results in up to four NF-186 molecules within a ∼190 nm spatial period (Fig. 4F and Fig. 6). Close-proximity NF-186 pairs, oriented in various directions along the AIS ∼52 nm spaced apart (Fig. 3C, E), were also observed as part of a cluster of NF-186 signal framed by Kv1.1 signal (Fig. 3A, F). We suggest that these pairs potentially represent two NF-186 molecules bound to a single AnkG molecule, consistent with the ∼52 nm cluster width of 3-4 gold particles detected by cryo-ET (Fig. 5F). The cluster size of two gold particles with a width of ∼29 nm abundantly present in cryo-ET data, likely reflects two secondary antibodies bound to one anti-NF-186 primary antibody. Such small clusters were not readily resolved in ExM due to the resolution limit of the setup. Although the nominal resolution after expansion is ∼27 nm, as described earlier, the effective resolution achieved in super-resolution microscopy experiments is typically lower due to linkage error, localization uncertainty and potential distortions introduced with expansion. Consistent with this, Airyscan imaging alone (without ExM) does not resolve the ∼190 nm spatial period of the MPS in our setup, despite a reported theoretical resolution of ∼120 nm at 488 nm excitation^40^. In this context, the ∼52 nm spacing of the closest NF-186 pairs approaches the practical resolving limit of our ExM experiments. At such near-limit separations, small positional uncertainties may occasionally prevent these pairs from being reliably distinguished, causing them to appear indistinguishable as a single structure.

The lack of strict alignment of NF-186 pairs with the K_v_1.1/AnkG organisation along the proximal-distal AIS axis (Fig. 4C) likely reflects antibody flexibility, the relative arrangement of NF-186 on AnkG, and potential conformational flexibility of the AnkG MBD. ExM (∼4.6x) revealed multi-molecule NF-186 organisation relative to AnkG but relied mainly on manual measurements. Complementary cryo-ET provided higher-resolution insight, exemplifying how the two approaches leverage each other’s strengths to uncover previously inaccessible aspects of the AIS architecture.

The data from cryo-ET (Fig. 5), similar to ExM, showing that NF-186 labelling appears in clusters or pairs of varying sizes, aligns with two AnkG molecules being situated in pairs of guided by the binding to β-IV-spectrin. The MBD of AnkG has been shown to harbour two possible binding sites for NF-186^47^, supporting our observation of close-proximity pairs potentially representing two NF-186 molecules bound to one AnkG and wider clusters representing more NF-186 molecules bound to two AnkG molecules on a single spectrin tetramer. The analysed clusters ranged from 2-16 gold particles, indicating that the AnkG MBD can bind several NF-186 copies or that AnkG pairs themselves are clustered axially. DBSCAN was performed by limiting clusters to a minimum of two particles with a maximum inter-particle distance of 54.4 nm. Occasional incidents of single particles may arise from unsaturated antibody labelling or from AnkG pairs harbouring one NF-186 alone and/or a pair of NF-186 and NrCAM, as these two cell-adhesion-molecules are shown to bind to the same part of the AnkG MBD^21^. Thus, single gold particles that did not belong to a cluster were excluded from the analysis (Supplementary Figure S7).

Altogether, our observations show that NF-186 organisation diverges from the conventional ∼190-nm periodic organisation of other AIS proteins (Fig. S6). This is consistent with earlier reports showing a large spread of NF-186 spacing in super-resolution microscopy imaging^9^.

These results show how immuno-gold labelling can support cryo-EM to study the ultrastructure of the AIS at a higher resolution than is achievable with super-resolution fluorescence microscopy. The analysis of the clustered NF-186 labelling in 2D projection images, shows a mean cluster-centroid distance of 199 ± 46.5 nm, confirming the overall periodic arrangement of membrane proteins in the AIS. Although, when analysing the labelling in 3D by cryo-ET, a more complex arrangement is revealed, as the labelling distribute at different Z-heights along the plasma membrane (Fig. 5E) and in clusters of varying size (Fig. 5F) matching the distances measured by ExM. Optimized labelling procedures will improve the possible output from this kind of analysis, as single molecule labelling is essential. *In situ* immunogold cryo-ET studies are currently limited to extracellular epitopes and would require alternative labelling strategies (e.g., nanobodies, genetically encoded tags) to access intracellular targets of intact cells. Regardless, imaging the labelled ultrastructure by cryo-ET allows for high resolution 3D analysis and could potentially improve the understanding of the arrangement of molecules within the general ∼190-nm spatial period of the AIS proteins. For one, cryo-ET is not limited in resolution in the Z direction unlike other super resolution methods.

Altogether, our results shed a light on the stoichiometry of AnkG-associated complex in the AIS which is a master organiser of this dense ultrastructure. Our study illustrates the possibilities of exploiting cryo-ET and ExM to move towards higher achievable resolution in visualization of the AnkG-associated complex. This approach may successfully address other questions such as the stoichiometry of Na_v_-AnkG molecules and the distance between the MBD and SBD of AnkG molecules. The u-ExM protocol used in this study resulted in ∼4.6-fold expansion of the samples which was sufficient to reveal new distances in the AIS and showing the applicability of this method in the study of AIS ultrastructure. ExM protocols ensuring larger expansion can be used to further push the effective resolution and unravel fine architectural details at an even smaller scale. Furthermore, both methods can visualize the periodic arrangement of the AIS in 3D at a higher resolution in the Z direction. This can provide insights into the axial organisation of the AnkG-associated complexes and the surrounding membrane proteins in the AIS in addition to the longitudinal periodic organisation. Finally, our study demonstrates the power of the collective use of the two methods, and that ExM was able to put the complex data from cryo-ET into perspective.

## Data Availability

The datasets generated and analysed during the current study are available from the corresponding author on reasonable request.

## Code Availability

Code is available at GitHub (https://github.com/jkdannerso). A snapshot of the code is archived at Zenodo (2D: DOI: 10.5281/zenodo.19068688 and 3D: DOI: 10.5281/zenodo.19068775).

## Acknowledgements

We thank the members of the H. B. Rasmussen (University of Copenhagen) and U. V. Nägerl (University of Bordeaux) laboratories, and M. F. Monreal from Bordeaux Imaging Center for helpful discussions on ExM analysis and valuable comments on the manuscript. We acknowledge Euro-BioImaging ERIC (https://ror.org/05d78xc36) for providing financial support to access imaging technologies and services via the French BioImaging-Node in Bordeaux, France. We are grateful to T. Boesen and the staff at the cryo-EM facility EMBION at Aarhus University for assistance with cryo-EM data acquisition and for helpful discussions on the analysis. We thank H. Dannersø for facilitating writing of the python programme for 2D-analysis of the gold clusters. We are grateful to C. Sun for valuable discussions.

This work was supported by a collaborative grant (BRAINSTRUC R155-2015-2666) and a professorship grant (R310-2018-3713) from the Lundbeck Foundation, infrastructure grants from the Danish Ministry for Research and Higher Education (EMBION - 5072-00025B) and the Novo Nordisk Foundation (NNF20OC0060483), equipment grants from the Carlsberg Foundation (CF22-1535 and CF23-1394), the CryoNet network grant supported by the Novo Nordisk Foundation, and a scientific exchange grant from EMBO to GB (9481).

## Author contributions

GB performed the fluorescence microscopy and ExM experiments and the relevant analysis. JKD performed the cryo-EM and cryo-ET experiments, created the script for 2D and 3D-cluster analysis and performed the relevant analysis. PN, UVN and LSL supervised the experiments. GB and JKD wrote the manuscript, SDSH, LSL, PN and UVN reviewed it. GB, JKD and SDSH created the figures presented in the manuscript.

## Supplementary Materials & Methods

### Supplementary Figures

**Figure S1.**
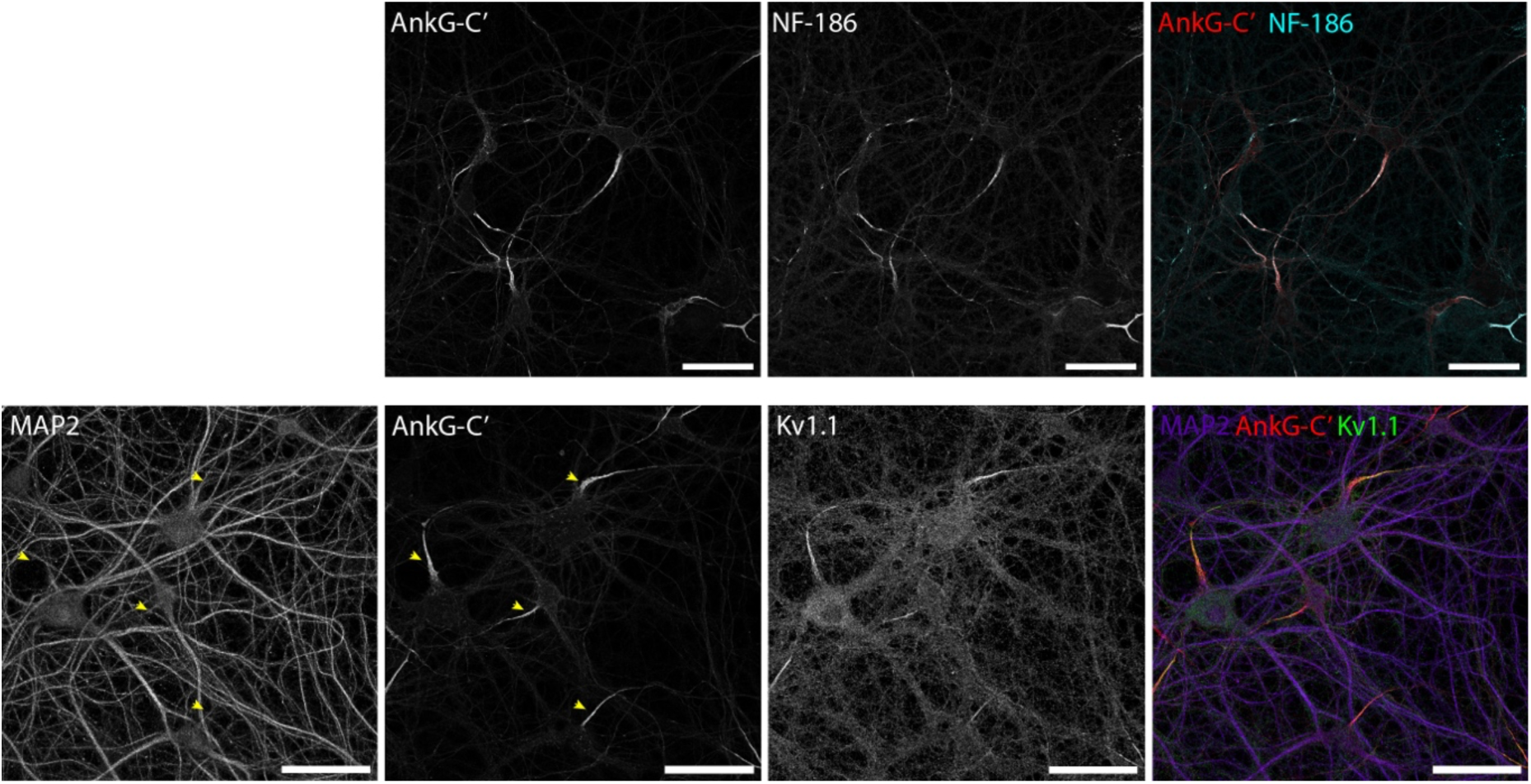
Confocal images of immunofluorescence staining in 19 DIV primary hippocampal neurons. Top, AnkG labelling using an antibody targeting the C-terminal tail (AnkG-C’; red) overlaps with NF-186 labelling (cyan; pan-Neurofascin antibody) in the AIS. Bottom, MAP2 labelling (purple) is absent from the AIS (yellow arrowhead), which is marked by AnkG-C’ and Kv1.1 (green) labelling. Scale bars, 50 µm.

**Figure S2.**
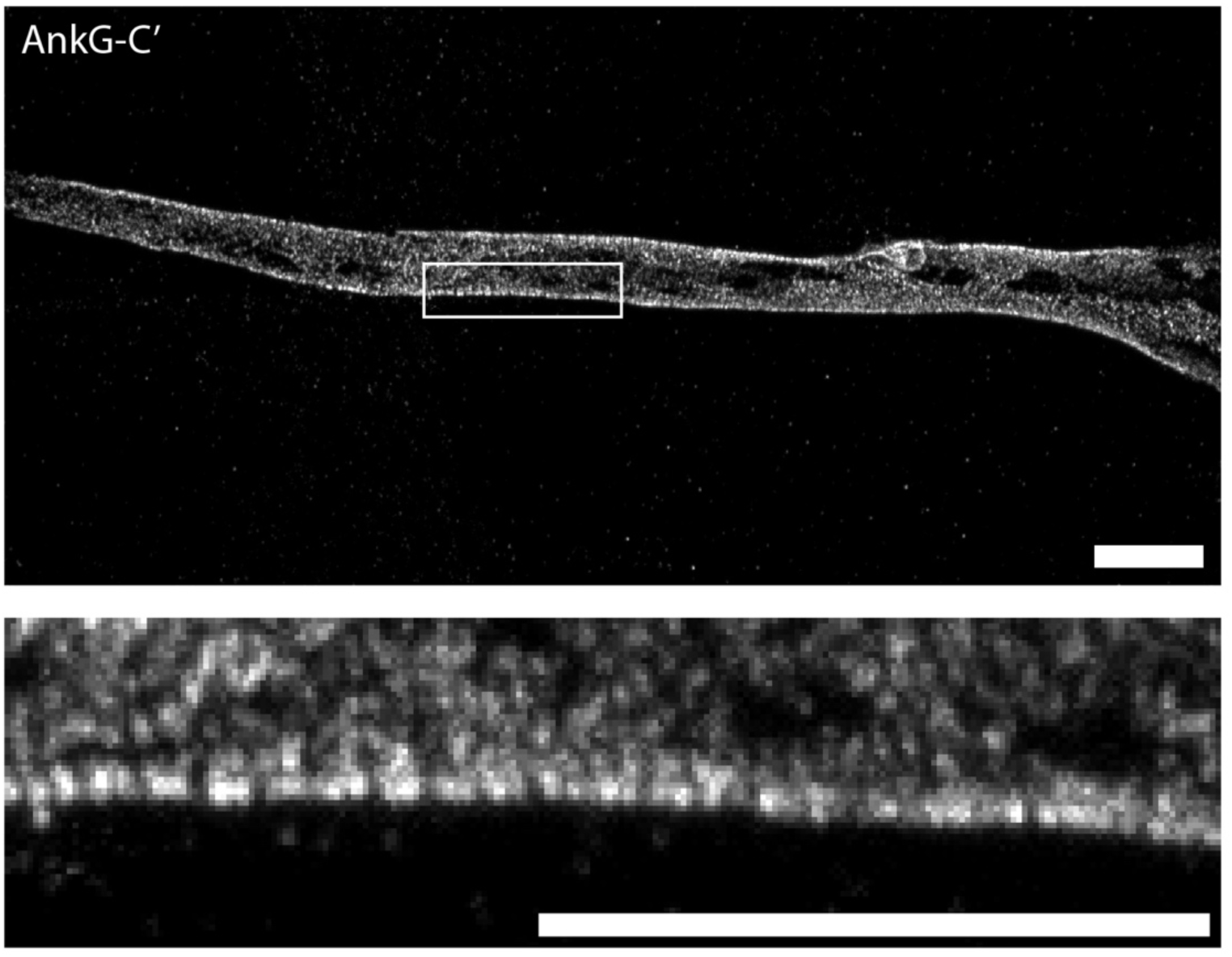
Confocal images of post-expansion AnkG-C’ labelling in the AIS. AnkG-C’ signal is observed in pairs. Maximum intensity projection of confocal images of expanded AIS. Scale bars, 10 µm post-expansion.

**Figure S3.**
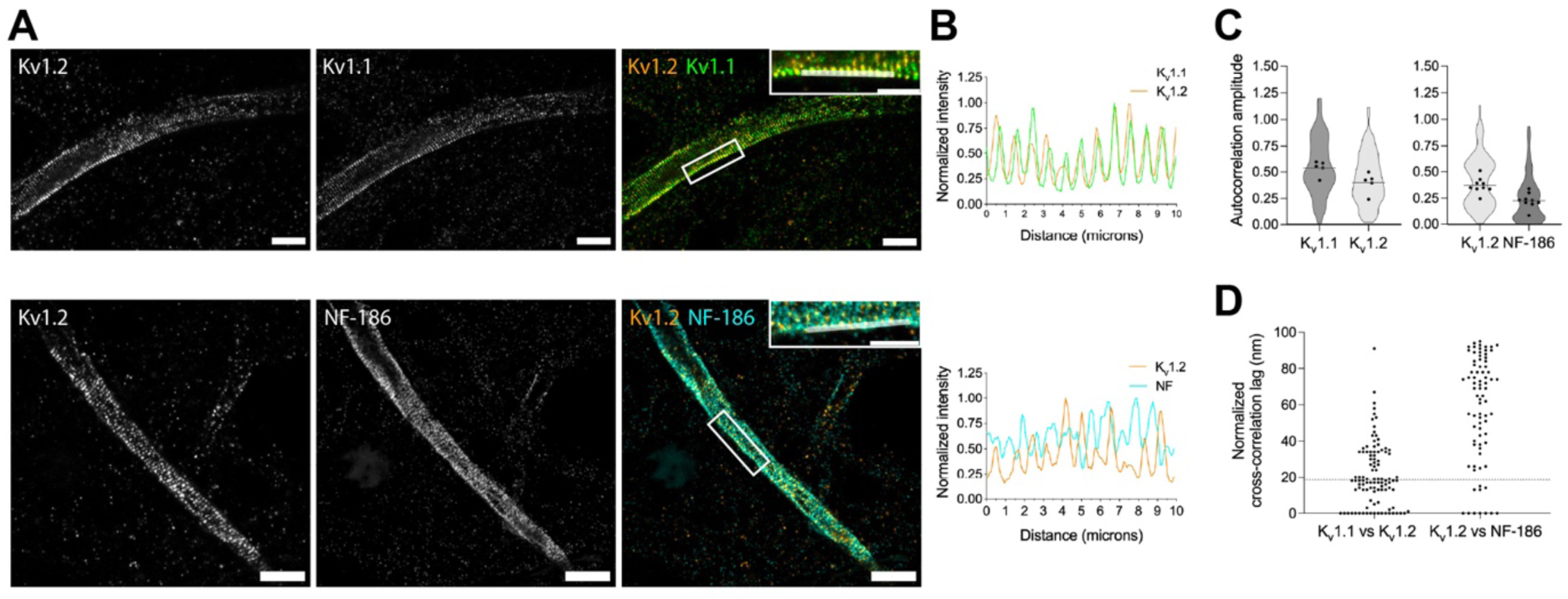
Pro-ExM resolves K_v_1.1 and K_v_1.2 colocalization interspersed with NF-186 in the AIS. **A,** Confocal images of ∼4.5x expanded AISs showing pre-expansion labelling of K_v_1.2 (orange) and K_v_1.1 (green), or K_v_1.2 and NF-186 (cyan) labelling. Close-ups show the white-framed regions in the overlayed images. Scale bars, 10 µm (full-frame images) and 5 µm (close-up images) post-expansion. **B,** Overlayed fluorescence intensity profiles along the white line ROIs in **A**. Top, in-phase K_v_1.2 and K_v_1.1 signals with ∼190 nm spacing. Bottom, out-of-phase K_v_1.2 and NF-186 signals. Distances are post-expansion. **C-D,** Independent cross-correlation analyses. **C,** Truncated violin plots show the autocorrelation amplitudes (AC) as an indicator of periodicity strength: K_v_1.1 (0.54 ± 0.07) vs K_v_1.2 (0.40 ± 0.10), and K_v_1.2 (0.37 ± 0.07) vs NF-186 (0.23 ± 0.07). AC: mean ± s.d., black dots: AIS-level means. **D,** Normalized cross-correlation lag in pre-expansion values. K_v_1.1 vs K_v_1.2 lag accumulates around 0-20 nm, while K_v_1.2 vs NF-186 lag accumulates around 75-95 nm, consistent with the in-phase and out-of-phase signals in panel **B**. Dashed line indicates the average pixel size after expansion factor correction (∼19 nm). Number of grids plotted: N = 102 from n = 5 AIS (K_v_1.1 vs K_v_1.2), N = 79 from n = 9 AIS (K_v_1.2 vs NF-186).

**Figure S4.**
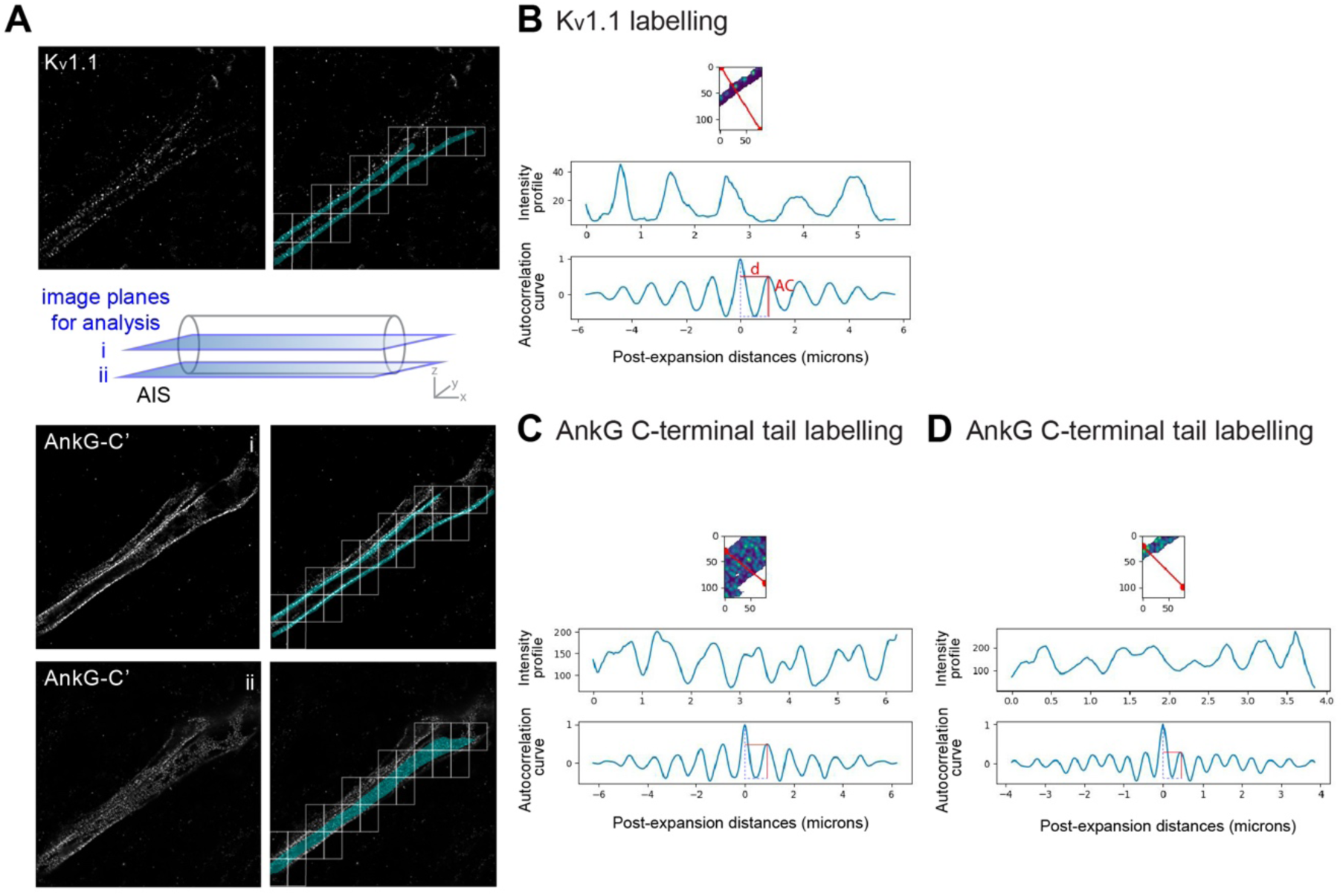
Autocorrelation analysis. **A,** Example fluorescence images of K_v_1.1 (top) and AnkG-C’ (bottom two images showing two Z planes) labelling in expanded AIS used for autocorrelation analysis (left column). Masks (blue) were applied to select labelled areas, and fixed rectangular grids (120×80 or 80×120 pixels) were generated for analysis (right column). The schematic illustrates the two Z planes (i, ii) of the AIS shown in the fluorescence images. **B-D,** Example autocorrelation grids of Kv1.1 **(B)** and AnkG-C’ **(C-D)** labelling. The red line on the grid indicates the direction of autocorrelation curve which is perpendicular to it. The respective intensity profiles and autocorrelation curves for the grids are depicted. Plots are generated using post-expansion distances. Autocorrelation amplitude (AC; the height of the first peak after the first valley) reflects the strength of the periodic pattern detected. The distance from the centre line to the first peak (d) shows the frequency of the periodic pattern, while the distance from the centre line to the first peak (d; frequency of the periodic pattern) represents inter-peak distance in the MPS. Mean inter-peak distances are similar in panels **B** and **C**, while panel **D** shows a shorter inter-peak distance.

**Figure S5.**
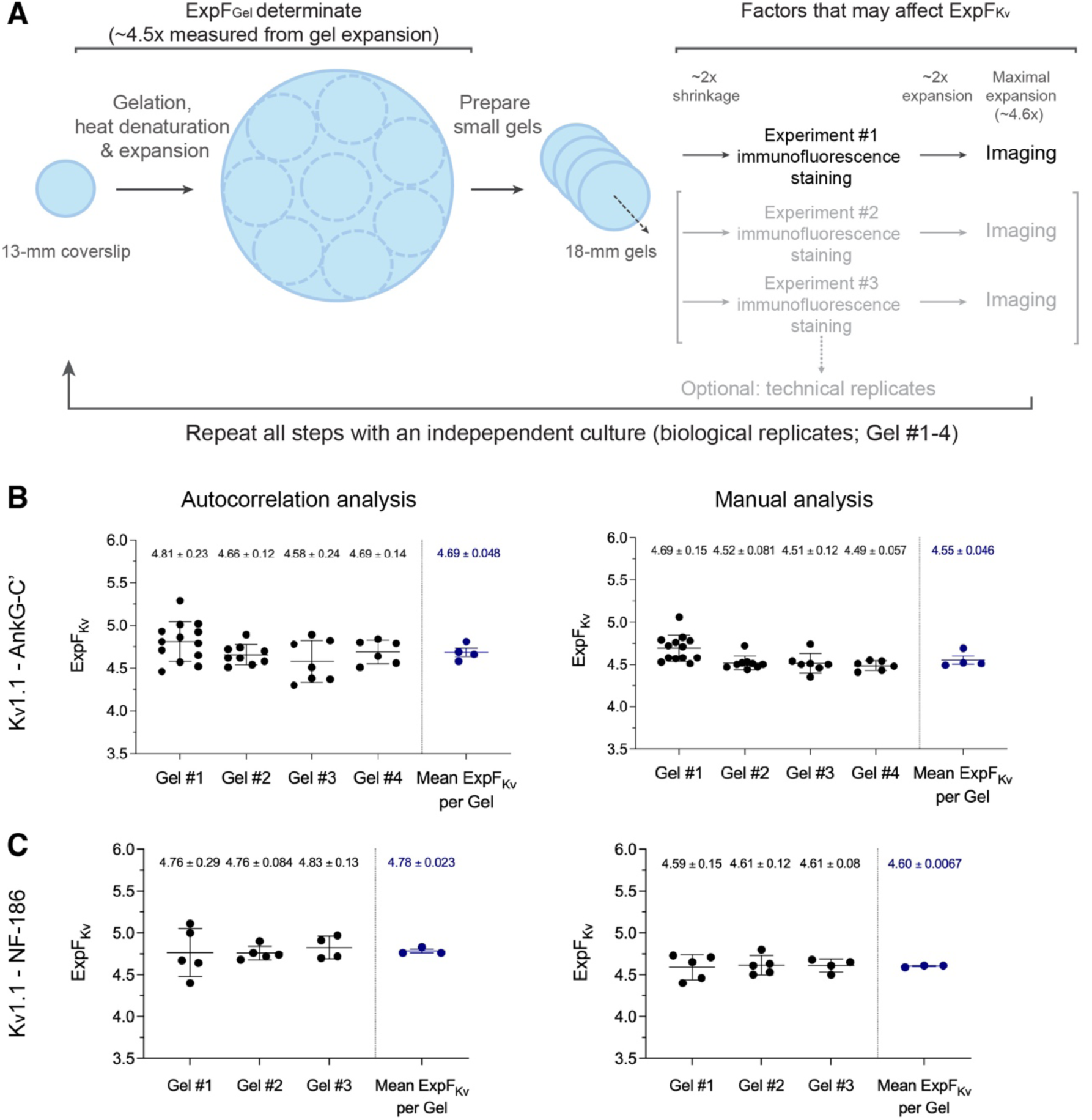
Schematics of the u-ExM experimental design and determination of expansion factor (ExpF). **A,** An initial expanded gel is prepared from a coverslip via gelation, heat denaturation, and expansion, defining the primary gel expansion factor (ExpF_Gel_). ExpF_Gel_ is calculated from the pre- and post-expansion dimensions of the gel as ∼4.5x in line with the u-ExM protocol by Gambarotto et al. (2021). The gel is then cut into smaller pieces (∼18 mm) for immunofluorescence staining. During staining, gels transiently shrink in PBS and re-expand in water prior to imaging, contributing to variability in the final expansion factor. To account for the expansion heterogeneity, a refined expansion factor (ExpF_Kv_) is calculated for each analysed AIS using the ∼190-nm spatial period of K_v_1.1 labelling as an internal reference. Multiple gel pieces from the same gel can optionally be used to repeat immunofluorescence staining as technical replicates. This procedure is performed across 3-4 independent cultures, serving as biological replicates (Gel #1, 2, 3, …). **B-C,** ExpF_Kv_ values calculated based on autocorrelation and manual analyses for gels with K_v_1.1 and AnkG-C’ (**B)** or K_v_1.1 and NF-186 (**C)** labelling. Black data points represent AIS-level mean ExpF_Kv_ values (mean ± s.d.), calculated from individual distance measurements. Blue data points represent gel-level mean ExpF_Kv_ values (mean ± s.d.), which were averaged to obtain a final ExpF_Kv_ value (mean ± s.e.m.). ExpF_Kv_ values vary across individual AIS, while mean ExpF_Kv_ values are more consistent across gels.

**Figure S6.**
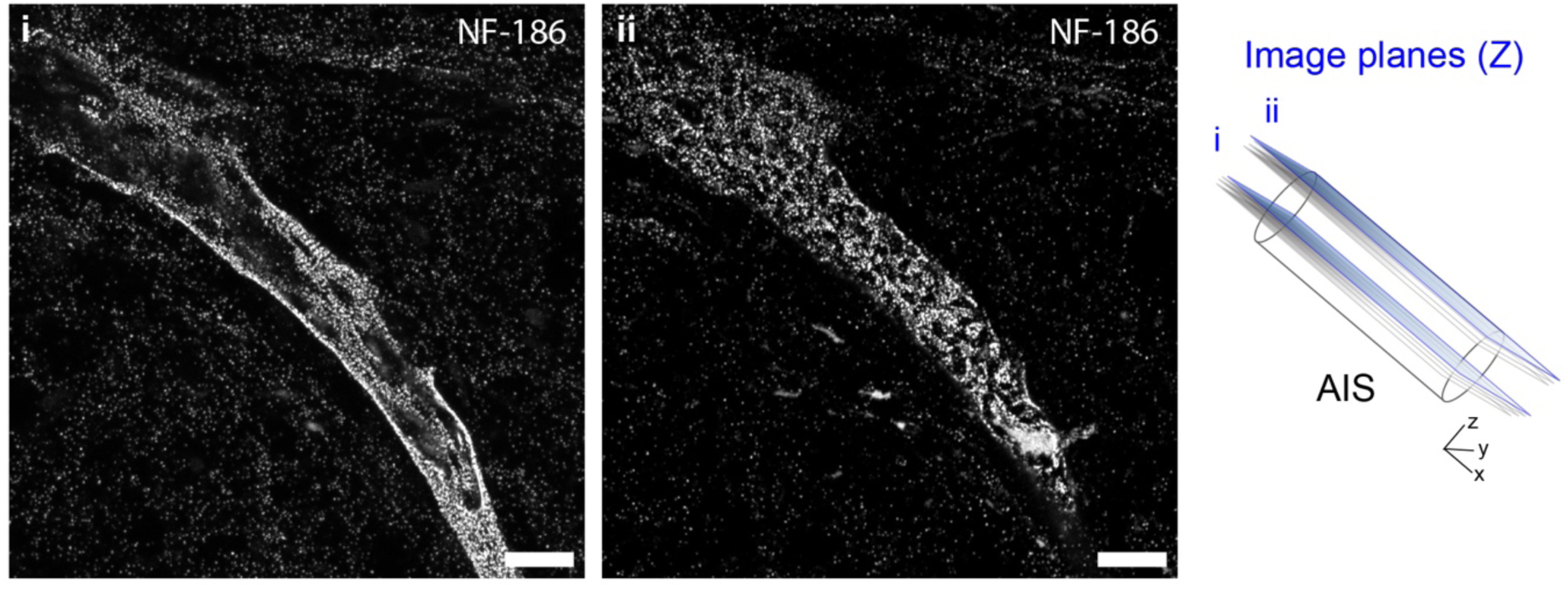
NF-186 labelling in the expanded AIS can be observed at various levels of periodicity. Maximum intensity projection of non-overlapping Z-stacks (i and ii) of the same confocal image of an expanded AIS with pre-expansion NF-186 labelling. NF-186 periodic organisation is more visible at a cross-sectional Z-plane (i) compared to a surface Z-plane (ii). Scale bars show 10 µm post-expansion.

**Figure S7.**
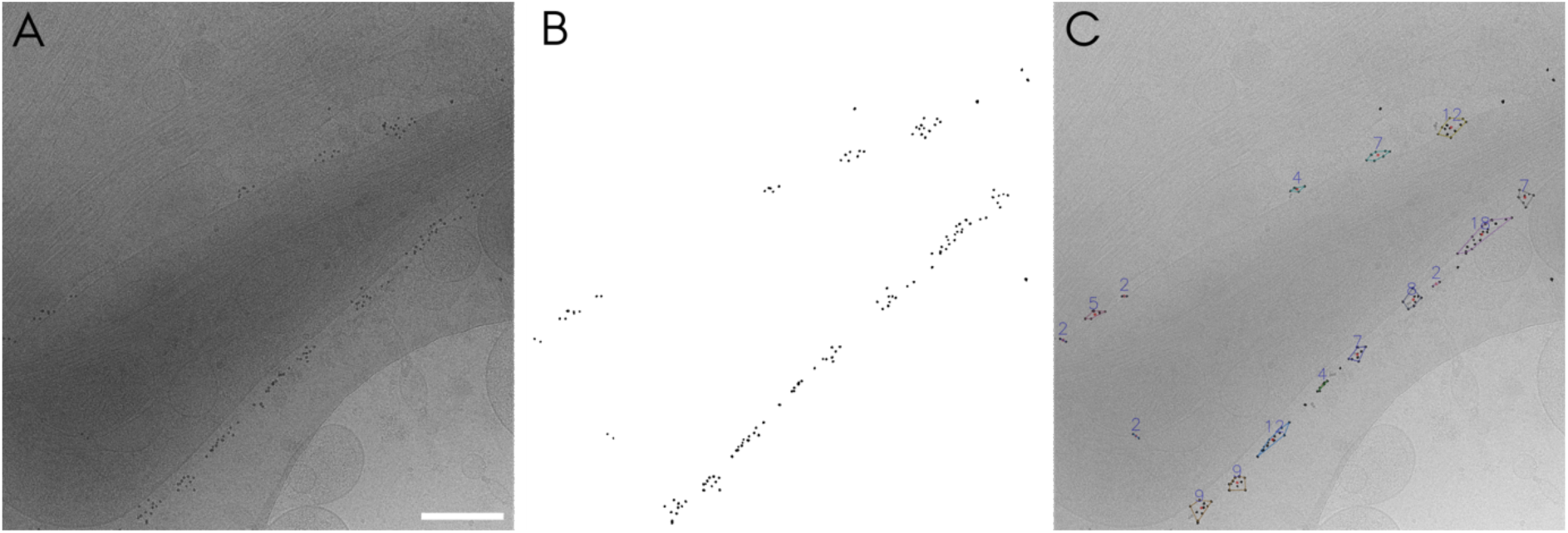
2-D Cluster analysis. **A,** Original projection image. **B,** The filtered image with only gold particles present. **C,** Overlay image with defined clusters of gold particles from which the manual analysis was performed by picking clusters in a row along the plasma membrane outputting inter-centroid distances.

**Figure S8.**
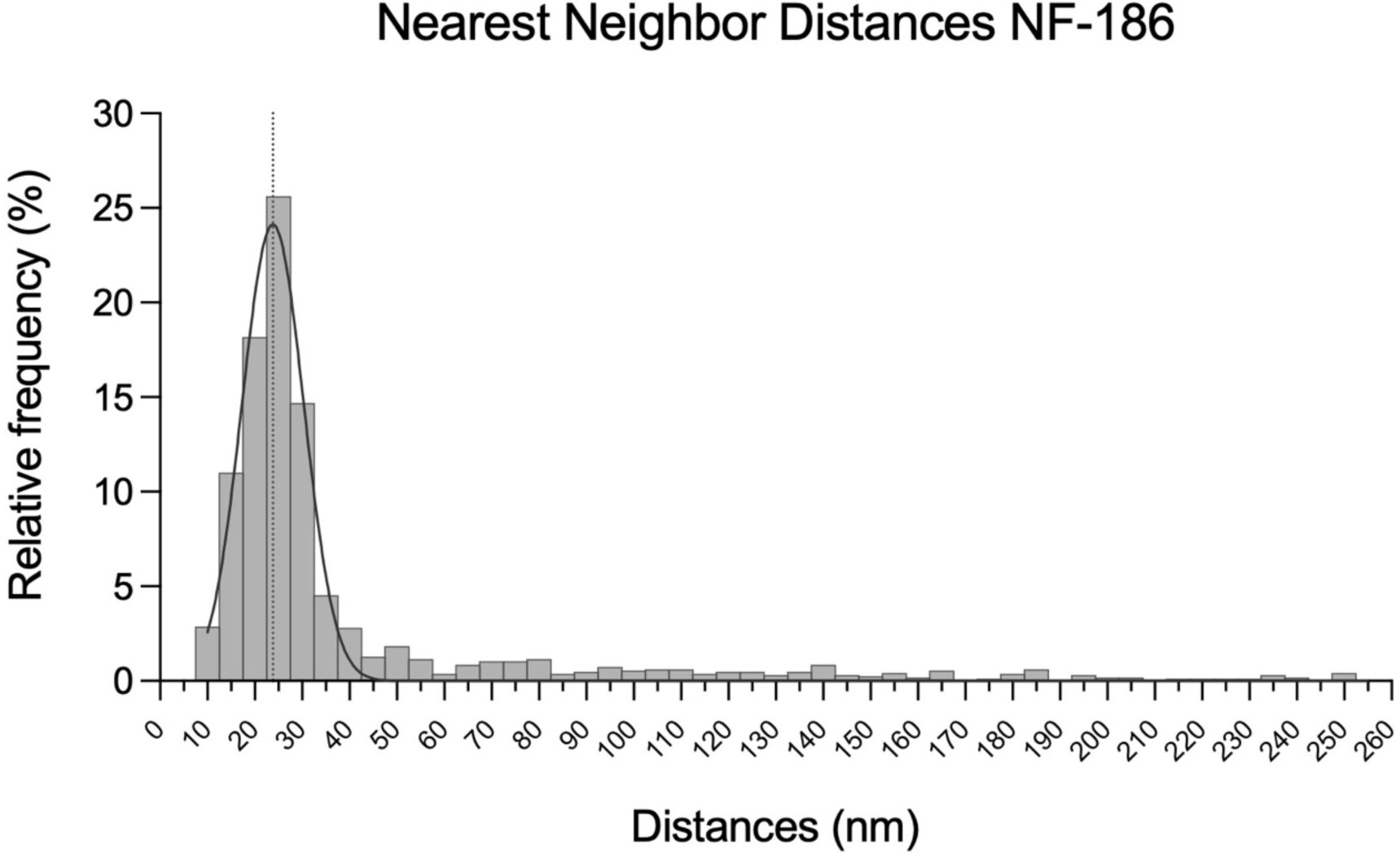
Nearest neighbour analysis of N=1636 points showing a peak at 23.8 nm. 1228 or 75% of the pairs have NN-distances < 35 nm.

### Supplementary Tables

**Table S1.** Primary antibody concentrations in immunofluorescence and immunogold labelling.

| **Antibody** | **Species** | **Brand, Catalog No.** | **Dilution**<br>**ExM** | **Dilution**<br>**ET** |
| --- | --- | --- | --- | --- |
| AnkyrinG C-terminal | Rabbit polyclonal | Synaptic Systems, 386 003 | 1:300 (u-ExM) | - |
| Pan-Neurofascin | Chicken polyclonal | Invitrogen, PA5-47468 | 1:70 (u-ExM)<br>1:20 (pro-ExM) | - |
| Pan-Neurofascin | Mouse monoclonal | NeuroMab, 75-172 | - | 1:100 |
| Kv1.1 (K36/15) | Mouse monoclonal | NeuroMab, 75-105 | 1:200 (u-ExM)<br>(5 µg/mL) | - |
| Kv1.1 (K36/15)<br>Supernatant | Mouse monoclonal | Developmental Studies<br>Hybridoma Bank,<br>K36/15 | 1:6 = 5 µg/mL<br>(u-ExM)<br>1:2.5 = 12 µg/mL<br>(pro-ExM) | - |
| Kv1.2 | Rabbit polyclonal | Abcam, ab123570 | 1:50 (pro-ExM) | - |

## Supplementary Methods

### Expansion Microscopy (ExM) with pre-expansion labelling

A pre-expansion immunolabelling protocol (pro-ExM) was followed as described by Asano et al. (2018) for cultured cells with minor changes. Briefly, 19 DIV neurons were fixed with 2% paraformaldehyde for 2 min at room temperature (RT) followed by ice-cold 100% methanol for 10 min at -20℃ and quenched in 100 mM glycine solution for 5 min at RT. Fixed neurons were permeabilized with 0.25% Triton-x100 in PBS for 15 min followed by blocking in 3% BSA in PBS for 1 h. Incubations with primary and secondary antibody solutions were done at overnight at 4℃ and for 2 h at RT, respectively. Primary antibodies used were listed on Supplementary Table S1. Donkey anti-chicken AlexaFluor 488 (1:100, Jackson Immuno Research 703-545-155), goat anti-mouse AlexaFluor 488 (1:500, Invitrogen A-11001) and goat anti-rabbit AlexaFluor 568 (1:500, Invitrogen A-11011) were used as secondary antibodies. Stained neurons were incubated in 0.1 mg/mL AcX solution for 3 h at RT. Gelation solution consisted of Stock X, Milli-Q water, Tetramethylethylenediamine (TEMED) and ammonium persulfate (APS) mixed at a ratio of 47:1:1:1. Coverslip was placed over a drop of 35 µL solution, cell side facing the solution, and was incubated for 1 h at 37℃ in the dark. The solidified gel attached to the coverslip was transferred into digestion buffer with 3 U/mL Proteinase K and incubated overnight at RT in the dark. The detached coverslip was removed, and the digestion buffer was replaced with Milli-Q water for allowing the gel expansion. The water was renewed two times every 30 min followed by 2 h expansion in the last round. A small piece was cut out from the expanded gel and mounted on a glass-bottom MatTek dish coated with PDL for imaging.

### Cross-correlation analysis

Cross-correlation analysis was performed using K2 Napari Wave Breaker plugin version 0.1.4^2^ integrated in Napari (doi:10.5281/zenodo.3555620) publicly available at https://github.com/SamKVs/napari-k2-WaveBreaker. The images to be analysed were treated the same way as in the autocorrelation analysis. In addition to autocorrelation amplitude and frequency, for each grid, a cross-correlation lag value was extracted at the angle and frequency where the two channels exhibited the highest average normalized autocorrelation. Cross-correlation lag describes the positional shift between the periodic pattern of the two analysed channels (i.e., K_v_1.1 and K_v_1.2 or K_v_1.2 and NF-186). The resulting lag values were then normalized to half of the 190 nm MPS unit (0–95 nm) to represent the positional shift, 0 nm describing full in-phase organisation, while 95 nm describing full out-of-phase organisation of the periodic pattern of the two analysed channels. The mean autocorrelation amplitude and frequency were calculated per AIS by averaging the values from only the grids used for cross-correlation lag, which were then averaged across all AIS.

